# Fast and reliable association discovery in large-scale microbiome studies and meta-analyses using PALM

**DOI:** 10.64898/2026.04.09.717497

**Authors:** Zhoujingpeng Wei, Qilin Hong, Guanhua Chen, Tina V. Hartert, Christian Rosas-Salazar, Suman R. Das, Meghan H. Shilts, Albert M. Levin, Zheng-Zheng Tang

## Abstract

Identifying microbial features associated with various covariates is a long-standing goal in microbiome research. Modern association studies incorporate an ever-increasing number of microbial features, covariates, and datasets from diverse cohorts. However, the complexity of microbiome data challenges analysis, often leading to poor replication of findings. We introduce PALM, a quasi-Poisson regression framework that enables fast and reliable association discovery in large-scale studies and meta-analyses. Extensive, realistic simulations demonstrate PALM’s advantages in controlling false discovery rates, boosting power, improving computational efficiency, and preserving cross-study homogeneity of association effects. Three real-world applications at different scales illustrate PALM’s utility, underscoring its potential to advance microbiome research.

## Background

Significant progress has been made in understanding the role of the human microbiome in health and disease^1,2^. Feature-level association analyses, commonly known as differential abundance analyses, are widely used to identify microbial features associated with clinical outcomes, host factors, and other relevant covariates. Recent advances have expanded microbial feature measurements beyond 16S rRNA gene amplicon profiling to multi-omics approaches, including metagenomics and metatranscriptomics^3,4^. Additionally, microbiome studies increasingly link microbial features to an ever-growing set of covariates^5,6^. For example, microbiome genome-wide association studies (mbGWAS) link microbial features to millions of host genetic variants. Furthermore, meta-analyses of microbiome association studies are becoming more common, aiming to enhance discovery power and identify microbial signatures that are generalizable across cohorts and populations^7–10^. The substantial growth in study scale offers an unprecedented opportunity to uncover novel microbial signatures, deepen our understanding of microbiome functions, and translate these insights into actionable interventions. However, poor replication of findings remains a significant barrier to achieving these goals. For instance, while existing mbGWAS have identified numerous genetic loci associated with microbial taxa, only a handful of the associations were shared across studies^11^.

Poor replication is due in part to the analytical challenges posed by the unique characteristics of microbiome data. Microbiome data consist of sequencing read counts mapped to microbial features. Because sequencing depth is uncorrelated with the actual microbial load in a sample, these counts are inherently compositional and are informative about relative abundances (RA) but not about absolute abundances (AA) per unit volume of the ecosystem^12,13^. These count data are inherently noisy. Measurement biases are introduced during experimental workflows (e.g., DNA extraction, primer binding, and amplification), leading to differential efficiencies in detecting features^14^. The majority of counts are concentrated in a few abundant features, with the remainder distributed among numerous low-abundance features. This results in a high proportion of zero counts, with the likelihood of observing zeros heavily influenced by a sample’s sequencing depth. Zero counts can also arise from true feature absence, which is difficult to distinguish from zeros due to insufficient sampling. The counts also exhibit substantial variation across samples, particularly when generated from different sequencing batches, study designs, or populations. These unique characteristics make microbiome data difficult to model.

Despite extensive research on microbiome association analysis, existing methods still struggle with inflated false discovery rates (FDR) and unstable effect estimation. In meta-analyses, two distinct forms of heterogeneity must be disentangled. Effect heterogeneity refers to genuine differences in the association parameter across studies, whereas distributional heterogeneity (batch effects) arises when observed microbiome profiles differ across studies because of technical or cohort factors (e.g., extraction protocols, sequencing platforms, or population structure)^9,15,16^. Batch effects alter the data distribution but do not, by themselves, imply that the underlying biological effects differ across studies. However, if study-level analyses are not properly specified, batch structure can propagate into the estimated effects, making truly homogeneous effects appear heterogeneous.

Connecting back to the unique characteristics of microbiome data, this problem can arise from several sources: (1) Data preprocessing. Many methods^5,17–19^ rely on normalization (sample-level scaling or transformations) and zero imputation; in meta-analyses, cross-study batch correction is also common^16^. These steps can distort target associations and propagate batch structure into association effects ^20–23^. (2) Compositionality. Many approaches^5,17^ target RA associations. Changes in one feature’s AA alter all RAs. Consequently, methods that ignore compositional effects can yield biased estimates of AA-level associations and propagate batch-induced shifts across features. Compositionality-aware methods^18,19,24^ improve FDR control but have not been comprehensively evaluated in meta-analysis settings. (3) Modeling and inference. Misspecified parametric models or unstable inference for sparse, over-dispersed counts can bias both effect estimates and their uncertainty quantification, leading to inflated FDR and apparent between-study effect heterogeneity in meta-analysis.

In this article, we introduce a semi-parametric quasi-Poisson method for Association analysis of Large-scale Microbiome studies and meta-analysis (PALM). PALM provides fast and reliable estimates of AA-level association effects with well-calibrated uncertainty quantification. In meta-analyses, PALM integrates the AA-level summary statistics across studies, preserving genuinely homogeneous effects and supporting reproducible, generalizable findings. We evaluate the performance of PALM through extensive realistic simulations for meta-analysis and illustrate its utility in three real data applications: identifying microbial features associated with colorectal cancer, hundreds of metabolites, and millions of host genetic variations. These studies show that PALM achieves superior FDR control, power, and speed and avoids spurious between-study heterogeneity in association effects.

## Results

### PALM overview

PALM is detailed in Methods, with a schematic overview for meta-analysis provided in Additional file 1: Fig. S1. In brief, PALM takes as input the microbiome count data, sequencing depth, the covariate of interest, and relevant confounders. For each study, PALM generates summary statistics for individual features, including the AA-level association effect estimates and their variances. These summary statistics are subsequently combined across studies to perform a meta-analysis and calculate overall association test *p*-values. The *p*-values are then adjusted for multiple testing over features, and a list of significant features exceeding the specified threshold is reported.

The modeling and estimation procedure of PALM offers a unique combination of advantages: (i) No data preprocessing requirements: PALM directly models count data, avoiding biases and instability introduced by data normalization, zero imputation, and batch effect correction. (ii) No distributional assumptions: PALM employs a semi-parametric approach that accommodates overdispersion in count data without relying on strict distributional assumptions. This results in more reliable inference compared to parametric models. (iii) Compositional effect correction: PALM connects association effects in the RA model to those in the latent AA model. Specifically, the RA-model intercept absorbs multiplicative nuisance from total microbial load and measurement bias, and RA effects equal AA effects minus a common, study-specific shift. By leveraging this relationship and the high dimensionality of features, PALM accurately corrects for compositional effects and recovers AA-level summary statistics. (iv) Fast and stable inference: PALM constructs summary statistics using score statistics, which improves numerical stability and accuracy for sparse and over-dispersed microbiome count data compared to Wald statistics. Importantly, these score statistics are computed by fitting a single null model independent of the covariate of interest, making PALM computationally efficient for association studies with large numbers of covariates. (v) Flexibility: PALM supports diverse study designs, such as cross-sectional and longitudinal studies, and allows for adjustment of confounders.

### Evaluations on realistic simulated data

We evaluated the performance of PALM and compared it with several leading methods in the meta-analysis of five studies testing for differential AA between two comparison groups. Each method generated summary statistics, which were combined using a fixed-effect meta-analysis, as the simulated association effects were homogeneous across studies. The compared methods included ANCOM-BC2^19^ and LinDA^18^, designed to detect differential AA. For ANCOM-BC2, we used the version that applies a feature-level sensitivity score filter to reduce potential false positives^19^. Therefore, only studies where a feature passed the sensitivity score contributed to the ANCOM-BC2 meta-analysis for that feature. Additionally, we included DESeq2^17^, MaAsLin2^5^, and LM-CLR^25^, which are widely used for detecting differential RA. LM-CLR is a linear model that uses the centered log-ratio (CLR)-transformed feature variable as the response.

We simulated microbiome data for five studies using five real metagenomics datasets as templates^8^. The real data exhibited systematic differences in the structures of microbial communities across studies (Additional file 1: Fig. S2a). To generate realistic simulated data, we used the microbiome data simulator MIDAsim^26^, which learned the structure of the template data for each study and produced data closely resembling the originals (Additional file 1: Fig. S3). Due to the information gap in realistically generating AA data from RA data, we directly introduced differential AA signals into the RA data and ensured the effects were homogeneous across studies (Methods). We evaluated a broad range of scenarios from the combinations of two sample size settings (*n* = 100, 120, 140, 160, 180 or *n* = 20, 40, 60, 80, 100), two feature size settings (*K* = 401 species or *K* = 92 genera), two association effect direction settings (balanced positive/negative effects or predominantly positive effects), and two sequencing depth settings (even or uneven depths between groups of comparison). For each scenario, we also varied the proportion of AA-differential features, ranging from 0.05 to 0.2.

We evaluated all methods in terms of FDR control and discovery power (Fig. 1). For all methods, we applied the Benjamini-Hochberg (BH) procedure^27^ for multiple testing correction and reported empirical FDR and power at a target FDR of 0.05. PALM was the only method that successfully controlled FDR across all scenarios. Unbalanced positive/negative effects and uneven sequencing depth exacerbated FDR inflation in the compared methods. PALM consistently maintained high power, even when the other methods exhibited FDR inflation. PALM exhibits the best power–FDR trade-off, as reflected by the area under the precision–recall curve (AUPRC; Additional file 1: Fig. S4).

**Fig. 1:**
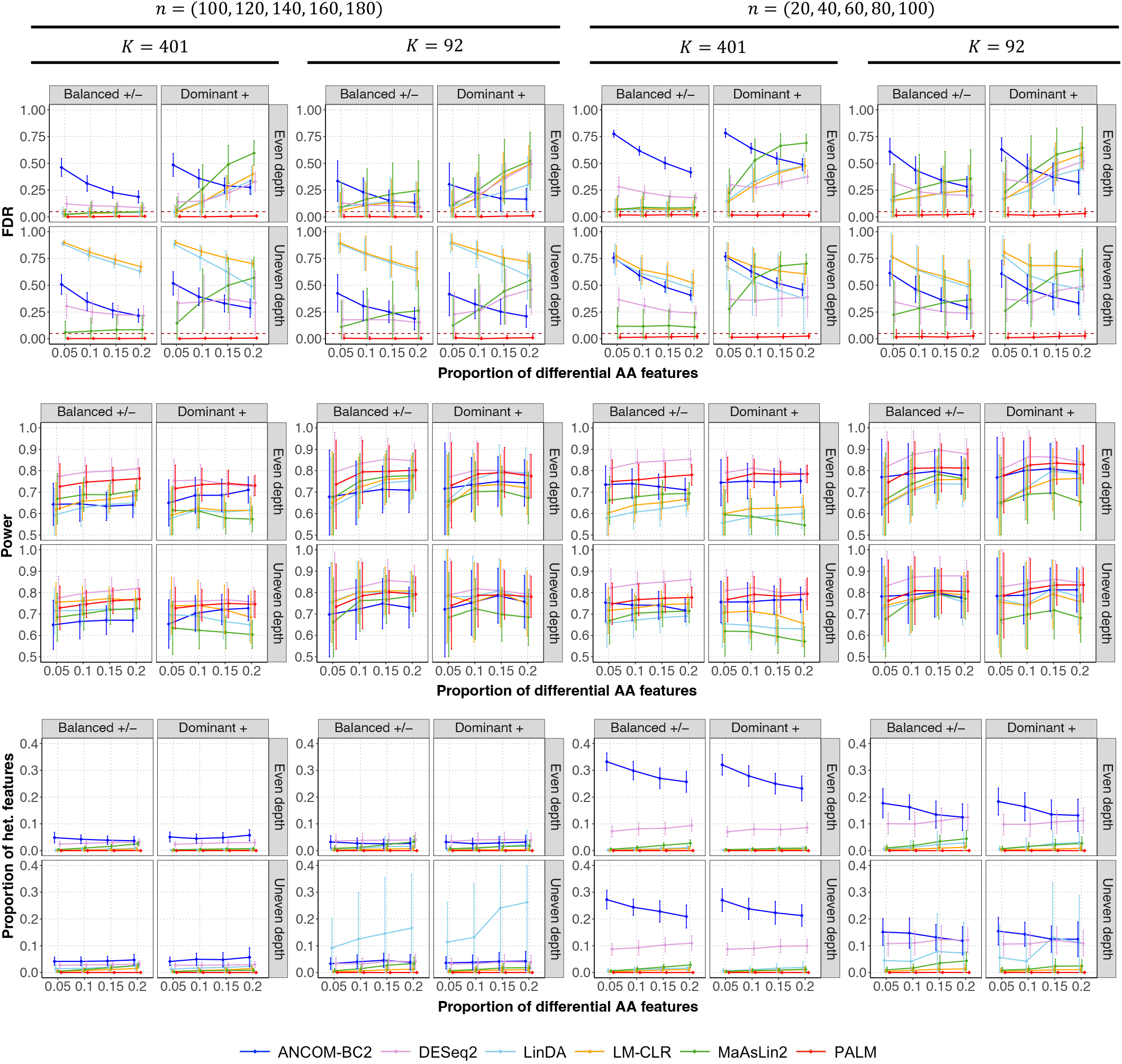
Comparison of different methods in the meta-analysis of five simulated studies with independent samples. The figure is divided into three sections, each representing a benchmark metric (y-axis): empirical FDR (top; the horizontal line indicates the target FDR), power (middle), and the proportion of heterogeneous features (bottom). Heterogeneous features are defined as those with variation in effect sizes across studies exceeding what is expected by chance (Cochran’s *Q* test *q*-value *<* 0.1). The x-axis represents the proportion of differential absolute abundance (AA) features. The top title shows the two scenarios of sample sizes *n* in five studies. The second title shows the two scenarios of feature size *K*. The column facet title represents two effect direction scenarios: Balanced +/−, where differential AA features have an equal probability of exhibiting positive or negative effects, and Dominant +, where all differential AA features exhibit positive effects. The row facet title represents two sequencing depth scenarios: one with even sequencing depth and one with uneven sequencing depth between the comparison groups. Each panel displays the mean estimated metrics with *±* standard errors (indicated by error bars) based on 100 simulation replicates.

Evaluating effect size heterogeneity across studies is crucial in meta-analysis as it provides key insights into the consistency and generalizability of study findings. To this end, we reported the proportion of features with significant heterogeneity (Fig. 1), determined using Cochran’s *Q* test (*q*-val *<* 0.1). In our simulations, the true AA-level effects were homogeneous; therefore, accurate summary statistics should exhibit no heterogeneity. PALM summary statistics showed no heterogeneity, indicating that it preserves genuinely homogeneous effects without introducing spurious differences. Other methods exhibited varying degrees of effect heterogeneity, with ANCOM-BC2 summary statistics showing the highest level in most scenarios.

We particularly evaluated method performance for rare features. In our simulation design, half of the AA-differential features were randomly drawn from relatively abundant features (average proportion ≥ 10^−3^ in the template data), and the other half from rare features (average proportion *<* 10^−3^). In the Additional file 1: Fig. S5, we reported results restricted to the subset of rare features. PALM maintains FDR control with good power, whereas DESeq2 and ANCOM-BC2 exhibit more pronounced FDR inflation than in the all-feature evaluation.

We also compared PALM meta-analysis with pooled-data analyses by other methods (Additional file 1: Fig. S6). Specifically, the original individual-level data from all studies were pooled together, and study indicators were adjusted as confounders in the association analysis. For ANCOM-BC2, the pooled-data analysis achieved better FDR control compared to its meta-analysis, but this came at the cost of reduced power. This trade-off occurred because the ANCOM-BC2 sensitivity score filter removed both false positives and true positives in the pooled-data analysis. For DESeq2, LinDA, LM-CLR, and MaAsLin2, the FDR inflation in the pooled-data analysis was similar to that observed in their meta-analysis.

Moving beyond studies with independent samples, we next evaluated the performance of methods in meta-analyzing studies with correlated samples (Fig. 2). For this evaluation, LM-CLR refers to the linear mixed model and DESeq2 is omitted because it cannot handle correlated samples. PALM effectively controlled the FDR, maintained high power, and exhibited no effect heterogeneity across studies. While the sensitivity score filter in ANCOM-BC2 effectively reduced FDR, it also lowered power. LinDA, LM-CLR, and MaAsLin2 exhibited FDR inflation and false effect heterogeneity across studies.

**Fig. 2:**
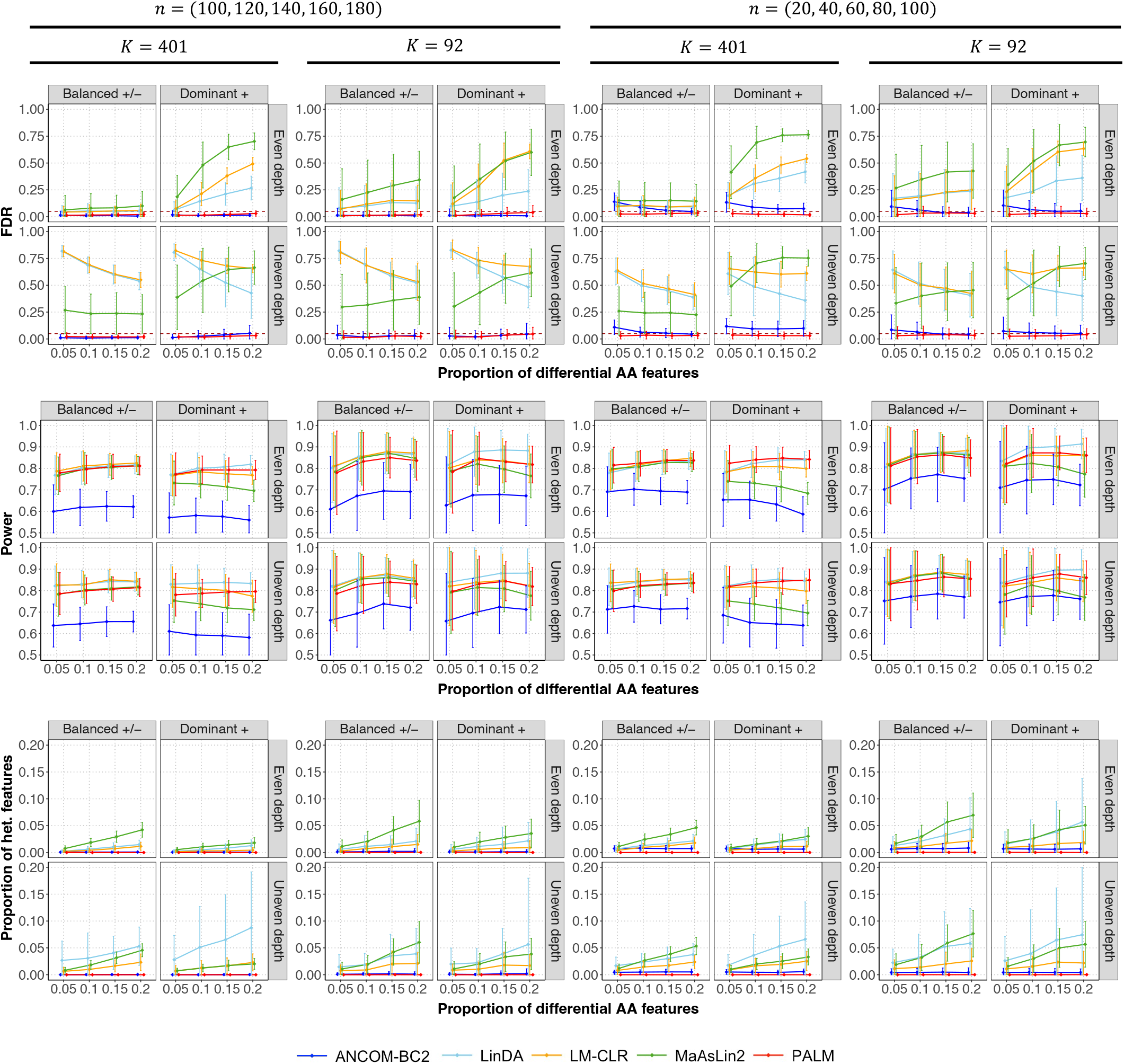
Comparison of different methods in the meta-analysis of five simulated studies with correlated samples. The figure is divided into three sections, each representing a benchmark metric (y-axis): empirical FDR (top; the horizontal line indicates the target FDR), power (middle), and the proportion of heterogeneous features (bottom). Heterogeneous features are defined as those with variation in effect sizes across studies exceeding what is expected by chance (Cochran’s *Q* test *q*-value *<* 0.1). The x-axis represents the proportion of differential absolute abundance (AA) features. The top title shows the two scenarios of sample sizes *n* in five studies. The second title shows the two scenarios of feature size *K*. The column facet title represents two effect direction scenarios: Balanced +/−, where differential AA features have an equal probability of exhibiting positive or negative effects, and Dominant +, where all differential AA features exhibit positive effects. The row facet title represents two sequencing depth scenarios: one with even sequencing depth and one with uneven sequencing depth between the comparison groups. Each panel displays the mean estimated metrics with *±* standard errors (indicated by error bars) based on 100 simulation replicates.

### Computation time

We quantified the computation time of different methods in meta-analysis across varying sample sizes and feature sizes as the number of covariates increased from 10^2^ to 10^5^ (Fig. 3). LinDA, LM-CLR, and PALM demonstrated the highest computational efficiency, making them suitable for large-scale association scans. Increasing the number of features and covariates led to longer computation times for all methods, but the impact was minimal for LM-CLR and PALM. DESeq2 and MaAsLin2 exhibited moderate computational efficiency, and ANCOM-BC2 had the highest computational demand. Overall, all three methods were computationally impractical for association scans with more than 10^5^ covariates.

**Fig. 3:**
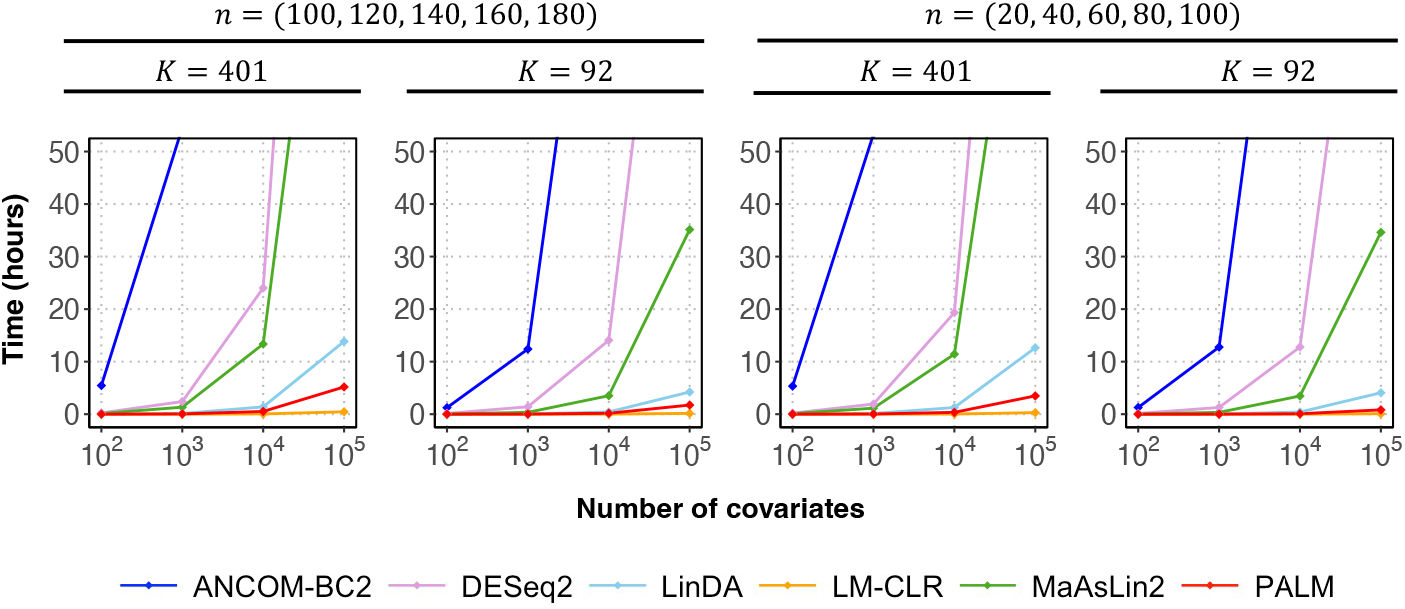
Comparison of computation time of different methods in the meta-analysis of five simulated studies with independent samples. The top title shows the two scenarios of sample sizes *n* in five studies. The second title shows the two scenarios of feature size *K*. Each plot shows the computation time in hours of different methods as a function of the number of covariates. The association scan was performed for one covariate at a time, and no parallel computing was used for all methods. Time beyond 50 hours is truncated. Values are averages over 100 simulation replicates for each setting.

### Meta-analysis of microbiome association studies for colorectal cancer

We applied PALM to three real data applications. In the first application, we meta-analyzed five metagenomics studies^8,28–31^ to identify gut microbial features associated with colorectal cancer (CRC). These five studies were conducted on three continents and differed in sampling procedures, sample storage, and DNA extraction protocols. The combined sample size across studies was 574, comprising 284 CRC cases and 290 controls (Additional file 1: Table S1). All raw sequencing data were reprocessed using a unified bioinformatics pipeline for taxonomic profiling, as previously described^8^. The microbial community structures across studies showed significant differences (PERMANOVA test *p*-value *<* 10^−5^, Additional file 1: Fig. S2a). After filtering out microbial species with less than 20% prevalence across all samples, 401 species were retained for analysis.

ANCOM-BC2, DESeq2, LinDA, LM-CLR, MaAsLin2, and PALM identified 117, 147, 120, 123, 126, and 84 species, respectively (Fig. 4a, Additional file 2: Table S2). A core set of 41 species was consistently identified by all methods, including well-known species associated with CRC, such as *Fusobacterium nucleatum, Bacteroides fragilis*, and *Parvimonas micra* ^32,33^. Notably, the majority of PALM discoveries (79%) were also identified by at least three other methods. In contrast, ANCOM-BC2 and DESeq2 respectively identified 27 and 34 species that were not supported by other methods. Most of these species were very low-abundance, with average proportions below 5 *×* 10^−4^. Furthermore, many of the DESeq2 discoveries (64) and ANCOM-BC2 discoveries (59) failed to pass the sensitivity score filter when ANCOM-BC2 was applied to pooled-data analysis, suggesting they may be potential false signals.

**Fig. 4:**
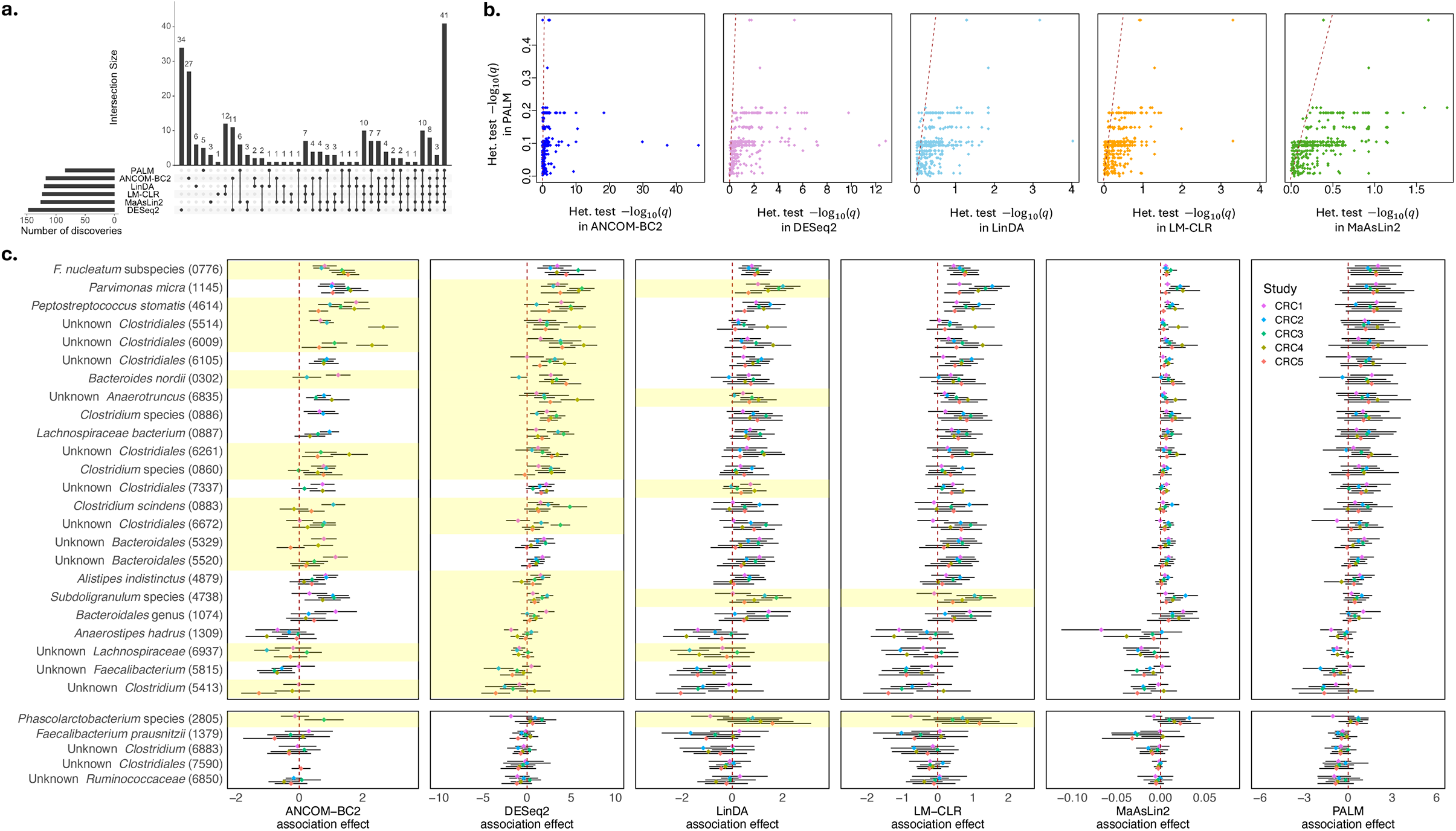
Results of the meta-analysis of microbiome association studies for colorectal cancer (CRC). **a**, The diagram shows the overlap of the CRC-associated species identified by different methods. **b**, Scatter plots of Cochran’s *Q* heterogeneity test − log_10_(*q*-value) in the meta-analysis using summary statistics of PALM (y-axis) and those of ANCOM-BC2, DESeq2, LinDA, LM-CLR, and MaAsLin2 (x-axis) over all species (dots). **c**, Upper panel: x-axis represents a core set of CRC-associated species identified by all methods but exhibiting significant heterogeneity in effects across studies in at least one method. Each box shows the summary statistics of a method, including the association effect estimates across studies (color-coded dots) and their 95% confidence intervals (black lines). In the box for ANCOM-BC2, the effects of some studies were not shown, as they were removed by the sensitive score filter. The row for species with significant heterogeneous effects across studies is shaded yellow in each panel (Cochran’s *Q* test *q*-value *<* 0.1); Lower panel: similar format as the upper panel but for the five species exclusively identified by PALM.

PALM exclusively identified five species, including *Faecalibacterium prausnitzii*, species under *Phascolarctobacterium*, and unknown members of *Ruminococcaceae, Clostridiales*, and *Clostridium. Faecalibacterium prausnitzii* and certain members of *Ruminococcaceae* are well-recognized as protective microbes against CRC^31,34^. These microbes contribute to gut health by producing short-chain fatty acids and promoting anti-inflammatory pathways that help maintain gut barrier integrity and modulate immune responses^35,36^. Recent research has also suggested that *Faecalibacterium prausnitzii* may enhance the effectiveness of immunotherapy for CRC while mitigating its adverse effects^37^. For these species, PALM effect estimates were consistently negative across studies, whereas effect estimates from the other methods showed both negative and positive associations (Fig. 4c, lower panel).

The summary statistics from different methods exhibited varying levels of cross-study effect heterogeneity. PALM summary statistics showed no effect heterogeneity across all species, whereas other methods displayed heterogeneity in some species (Cochran’s *Q* test *q*-value *<* 0.1), with the strongest heterogeneity observed in ANCOM-BC2 and DESeq2 summary statistics (Fig. 4b, c). While it is impossible to know whether true effect sizes are heterogeneous in real data, our simulation results suggested that high observed heterogeneity may arise from inaccurate effect inference, even when the true effect sizes are homogeneous.

### Meta-analysis of microbiome studies linking to the metabolome

In the second application, we conducted a meta-analysis of eight microbiome-metabolome association studies^15,38–44^ to identify gut microbial features associated with gut metabolites. The total sample size across all studies was 2,127. The studies differed in microbiome sequencing platforms, metabolomic measurement methods, cohort demographics, and study designs (Additional file 1: Table S3). All raw sequencing data were reprocessed and metabolite identifiers were consolidated for uniformity, as described elsewhere^45^. The microbial community structures across studies showed significant differences (PERMANOVA test *p*-value *<* 10^−5^, Additional file 1: Fig. S2b). Most metabolites were quantified in only a subset of studies. To ensure better data quality, we focused on metabolites and genera with at least 10% prevalence within individual studies and excluded metabolites present in only one study. The final datasets for analysis contained 450 metabolites and 101 genera, yielding 43,722 metabolite-genus pairs for the association scan. Summary statistics for individual studies were generated by treating each metabolite as a covariate of interest and adjusting for disease status as a confounder.

LinDA, LM-CLR, and MaAsLin2 identified significantly more features than PALM across metabolites (Fig. 5a, Additional file 3: Table S4). While ANCOM-BC2 identified a comparable number of features to PALM (Fig. 5a), the overlap between features identified by the two methods was low (Additional file 1: Fig. S7). In addition, we evaluated the number of associated metabolites across all features and characterized the top features associated with a large number of metabolites in the results of each method (Additional file 1: Fig. S8). Compared to other methods, the top features identified by PALM tend to be more abundant across individuals and are considered part of the human core microbiota^46,47^. We also found that this trend was driven by positive associations rather than negative associations (Additional file 1: Fig. S8b,c). Indeed, many of these features, such as *Phocaeicola, Bacteroides, Anaerostipes, Blautia, Faecalibacterium*, and *Roseburia*, are well-known producers of various metabolites, particularly short-chain fatty acids and other bioactive compounds essential for host health^48–50^.

**Fig. 5:**
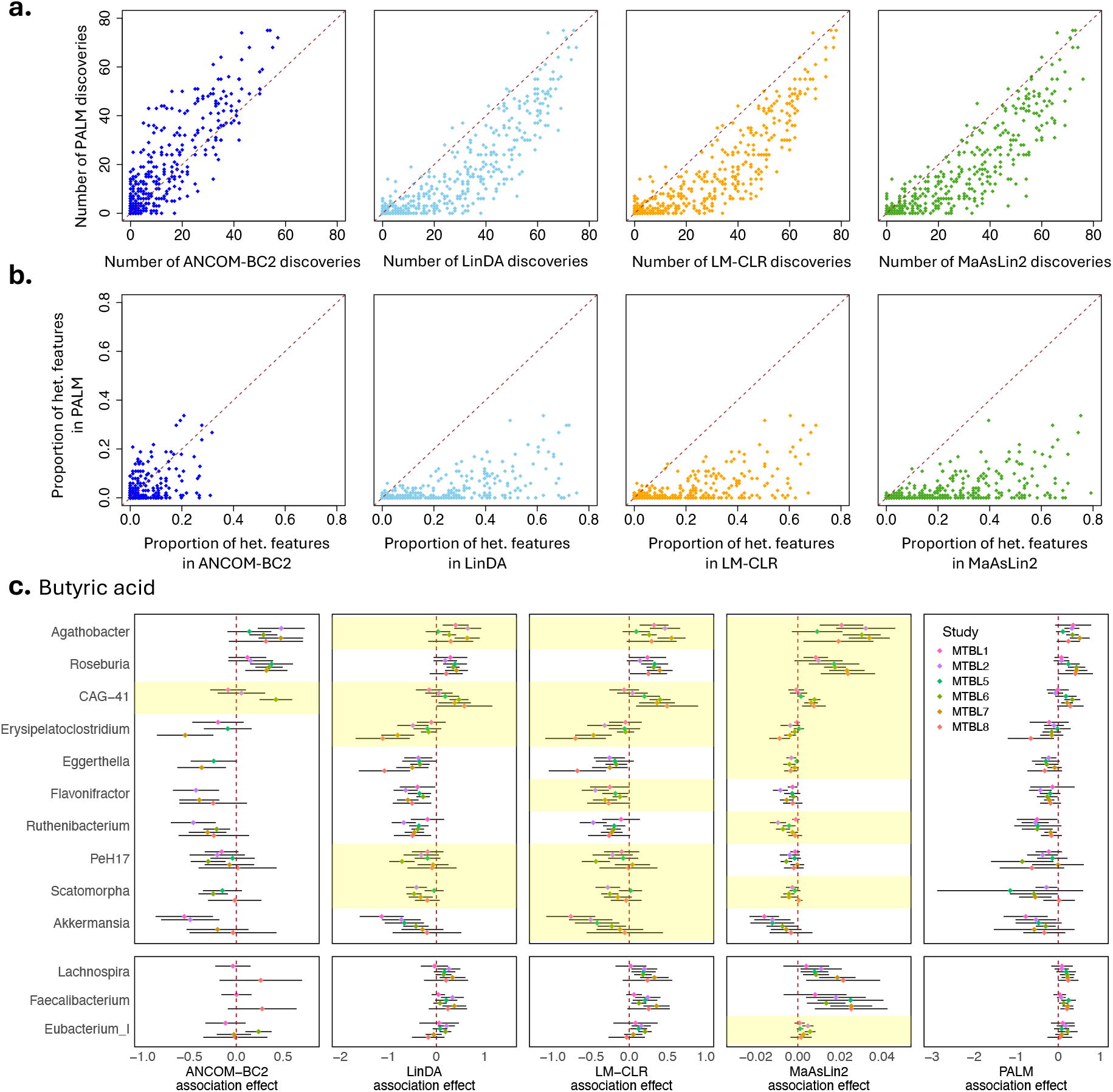
Results of the meta-analysis of microbiome-metabolome (MTBL) association studies. **a**, Scatter plots of the number of features identified by PALM (y-axis) and by ANCOM-BC2, LinDA, LM-CLR, and MaAsLin2 (x-axis) over all metabolites (dots). **b**, Scatter plots of proportions of features with significant cross-study effect heterogeneity (Cochran’s *Q* test *q*-value *<* 0.1) based on summary statistics of PALM (y-axis) and ANCOM-BC2, LinDA, LM-CLR, and MaAsLin2 (x-axis) over all metabolites (dots). **c**, Upper panel: x-axis represents a core set of butyric-acid-associated features identified by all methods but exhibiting significant heterogeneity in effects across studies in at least one method. Each box shows the summary statistics of a method, including the association effect estimates across studies (color-coded dots) and their 95% confidence intervals (black lines). In the box for ANCOM-BC2, the effects of some studies were not shown, as they were removed by the sensitive score filter. The row for features with significant heterogeneous effects across studies is shaded yellow in each panel (Cochran’s *Q* test *q*-value *<* 0.1); Lower panel: similar format as the upper panel but for the three well-known butyric-acid-associated features not included in the upper panel.

We further investigated cross-study effect heterogeneity in the summary statistics generated by different methods. For almost all metabolites, LinDA, LM-CLR, and MaAsLin2 identified a larger proportion of features with heterogeneous effects compared to PALM (Fig. 5b). As an example, we highlighted the association results for butyric acid (Fig. 5c; additional results for cholic acid are shown in Additional file 1: Fig. S9). Many features associated with butyrate exhibited significant effect heterogeneity in the summary statistics produced by LinDA, LM-CLR, and MaAsLin2. The lower panel of Fig. 5c presents results for three well-known taxa contributing to butyrate production. These taxa were identified by LinDA, LM-CLR, MaAsLin2, and PALM but were missed by ANCOM-BC2. Their PALM effect estimates were consistently positive across studies, but the ANCOM-BC2 effect estimates had inconsistent directions.

### Analysis of microbiome and host genetics associations

In the third application, we conducted a gut microbiome and host genetics association study in a single infant cohort. The cohort consisted of 502 healthy term infants, with their gut microbiomes measured once during their first year of life. In our analysis, the microbial features included 109 amplicon sequence variants (ASVs) with a prevalence greater than 5%, and the covariates comprised 6,174,920 SNPs with a minor allele frequency (MAF) *>* 5%, resulting in about 6.7 *×* 10^8^ ASV-SNP pairs for the association scan. We applied computationally feasible methods LinDA, LM-CLR, and PALM to perform the mbGWAS. Several existing mbGWAS studies have also used LM-CLR in their analyses^25,51^. In all methods, we adjusted for age at enrollment, sex, the top five ancestry principal components, delivery mode, antibiotic exposure, and birth season as confounders.

Our analyses were completed within 20 hours using 48 parallel cores on a 2.30 GHz Intel Xeon Gold 5118 processor. Using the genome-wide significance threshold (*p*-value *<* 5 *×* 10^−8^), LinDA identified 71 ASVs associated with at least one SNP, LM-CLR identified 50 ASVs, and PALM identified a single ASV (ASV2) associated with three SNPs (the top SNP is rs7780824 with *p*-value = 6.4 *×* 10^−9^, Fig. 6a). Many ASVs associated with a large number of SNPs in the LinDA and LM-CLR results tend to have low abundance (Additional file 1: Fig. S10a). Both methods rely on the CLR transformation, which requires adding a pseudo-count to handle zeros. To assess the impact of the pseudo-count, we re-ran LinDA and LM-CLR, varying the pseudo-count from the default value of 0.5 to 0.01. While both methods still identified many significant hits (Additional file 1: Fig. S10b), the majority differed from those identified previously (Fig. 6b), suggesting that the findings in LinDA and LM-CLR were unstable and may include false positives. The PALM-identified ASV2 belongs to genus *Escherichia-Shigella* and is highly prevalent in the cohort (92%), which naturally enhances its statistical power for detection in a moderate-sized GWAS. A clear trend of increased average relative abundance of ASV2 is observed in individuals with higher alternative allele dosages for rs7780824 (Fig. 6c).

**Fig. 6:**
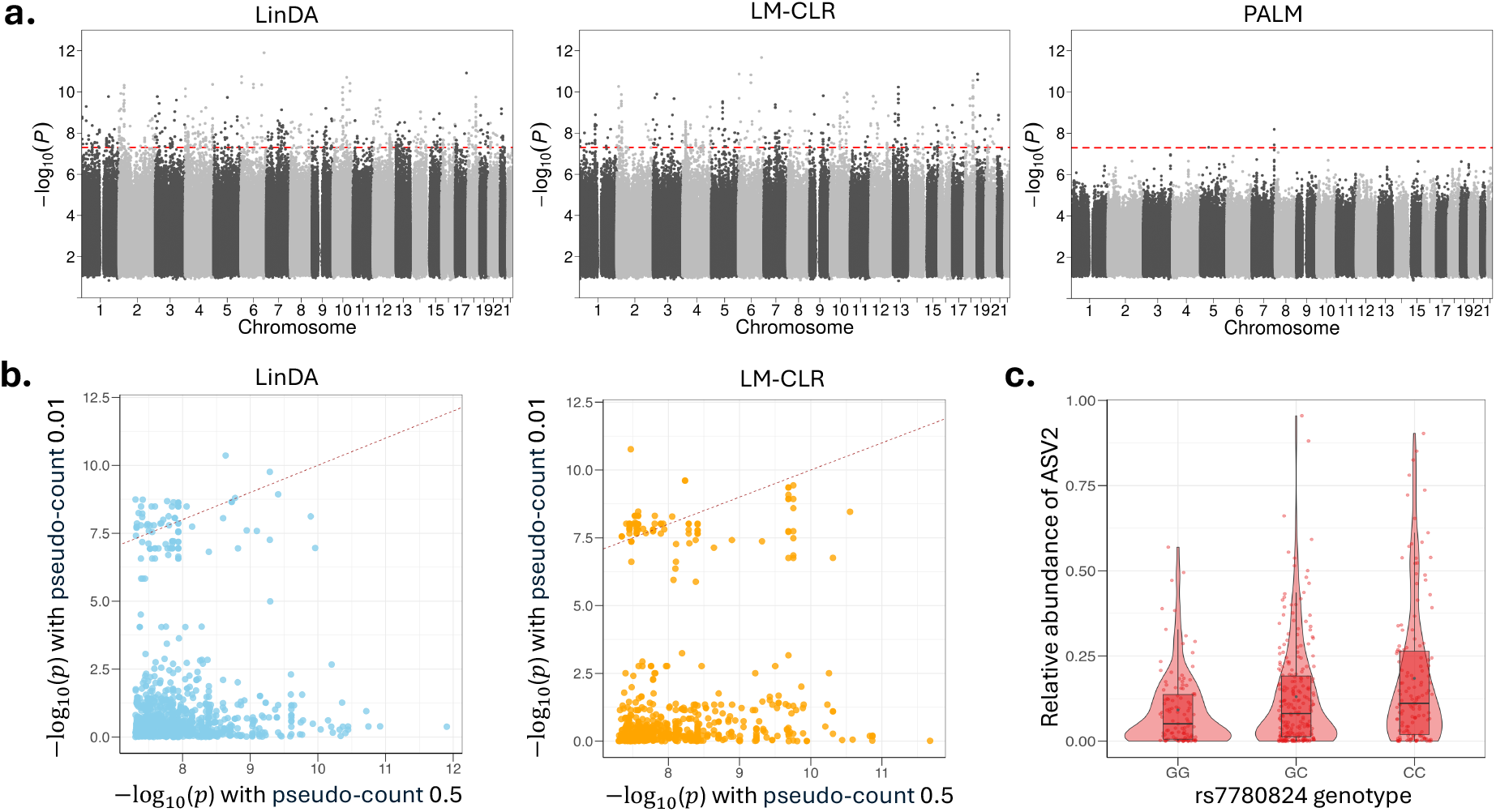
Results of the microbiome and host genetics GWAS. **a**, Manhattan plot aggregating the top associations with the microbial feature. Each SNP was tested against each of the 109 ASVs and the Manhattan plot shows the lowest resulting *p*-value for each SNP. The horizontal dashed line marks the genome-wide significance (*p*-value = 5 *×* 10^−8^). The top hit in PALM results involves feature ASV2 and SNP rs7780824. **b**, Comparison of *p*-values when using the pseudo-count of 0.5 (default) and 0.01 in LinDA and LM-CLR analysis. Each dot represents a significant ASV-SNP pair in the analysis using the default pseudo-count. **c**, Relative abundances of ASV2 stratified by groups of individuals based on the rs7780824 genotype. Data points are represented as dots. The boxplots show the median (center line), the first and third quartiles (box edges), and whiskers extending to 1.5 times the interquartile range. Violin plots illustrate the density distribution of the data points.

## Discussion

The unique characteristics of microbiome data pose significant analytical challenges, which contribute to the poor replication of findings, particularly in large-scale association studies and meta-analyses. We introduce PALM, a statistical framework designed for fast and reliable association discovery. PALM directly models feature counts using quasi-Poisson regression to infer AA-level associations without requiring data preprocessing or distributional assumptions. By leveraging score statistics, PALM achieves computational efficiency and reliable association effect inference. Furthermore, PALM preserves homogeneous association signals across studies, even when microbial communities exhibit strong batch effects due to differences in technical batches, populations, study designs, and other factors.

We conducted comprehensive and realistic simulation studies to validate the performance of PALM in the meta-analysis of microbiome association studies. Recent research highlights that unrealistic simulations can compromise the fair benchmarking of microbiome differential abundance methods^52^, underscoring the importance of assessing the similarity between simulated and real data. In our simulations, we used five real microbiome datasets as templates to generate realistic multi-cohort microbiome datasets. The simulated data closely mirrored the structure of the real data (Additional file 1: Fig. S3), providing strong support for the reliability of our method comparisons. The simulations showed that PALM was the only method to consistently control FDR over diverse scenarios. It also demonstrated high power in identifying true covariate-associated features at the AA level. In our simulations, where the introduced association effects were truly homogeneous across studies, PALM was the only method to provide homogeneous association effect estimates over all scenarios. The performance of PALM in real data applications aligns with these simulation results. In the meta-analyses, PALM summary statistics exhibited lower cross-study heterogeneity compared to those of other methods. While PALM generally identified fewer microbial features than other methods, its discoveries were more stable and biologically meaningful.

To estimate the common compositional shift in RA effects, we use the median of the estimated RA effects. This estimator performs well in our simulations with up to 20% of differential AA features. We also examined a more challenging setting with 40% differential AA features (Additional file 1: Fig. S11). In this case, compositional bias is larger (especially when all effects are positive) and all methods show more severe FDR inflation; although PALM with the median compositional bias correction continues to yield lower empirical FDR than the other methods, it no longer controls FDR at the nominal level when all effects are positive. To address this, we propose an alternative estimator of the compositional effect (Additional file 1: Supplementary Methods 1.3). This approach adapts the idea in the microbiome meta-analysis method Melody^53^: we treat study-specific compositional biases as hyperparameters in a summary-statistics–based best-subset selection framework and jointly tune these correction factors across studies together with the subset-size hyperparameter. This approach is slower than using the median directly, and when the proportion of differential AA features is small, as in the main simulations, its effectiveness at correcting compositional bias is similar to that of the median (Additional file 1: Fig. S12).

The design of PALM score-statistic-based test enables it to efficiently handle association analyses involving a large number of features and covariates. Unlike many existing methods, it requires fitting a null model only once, regardless of the number of covariates, making it highly computationally scalable. This advantage was demonstrated in the mbGWAS application, where PALM efficiently analyzed millions of genetic variants. We also evaluated the Wald test under the PALM model. Although it attains higher power, it exhibits greater FDR inflation, particularly with small sample sizes (Additional file 1: Fig. S13). The score-statistic–based test demonstrates a more favorable power–FDR trade-off than the Wald test (Additional file 1: Fig. S14).

The current implementation of PALM is based on the quasi-Poisson model, which assumes a linear relationship between the variance and the mean. However, this mean-variance relationship may not hold for all microbial features. The PALM framework could be extended to incorporate the quasi-gamma-Poisson model, allowing the variance to follow a quadratic relationship with the mean. This article focuses on testing associations with one covariate at a time. Extending PALM to support multi-covariate association analyses would broaden its utility and application scope.

### Conclusions

High-throughput sequencing technology empowers researchers to profile microbiomes, yet the resulting data are highly complex, capturing only the noisy relative abundances of various microbial features. PALM addresses critical methodological gaps in modern large-scale microbiome association studies by integrating robust statistical modeling, computational scalability, and effective meta-analysis capabilities. By delivering reliable and generalizable results, PALM will accelerate discoveries with translational potential and advance microbiome research in health and disease. Beyond microbiome studies, PALM can be readily adapted to other types of high-dimensional compositional omics data, such as metabolomics and proteomics. We have provided an efficient R package for the broad utility of the method.

## Methods

### PALM framework

We first elucidate the relationship between RA and AA associations, laying the foundation for PALM’s approach to recovering AA associations. We then introduce the PALM framework in the context of meta-analysis.

#### Relationship between AA and RA associations

Consider *n* independent samples and *K* microbial features. For sample *i*, let *X*_*i*_ be the covariate of interest, *N*_*i*_ the sequencing depth, *Y*_*ik*_ the read count assigned to feature *k, W*_*ik*_ the (latent) AA of feature *k* in a unit volume, and 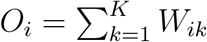 the total microbial load. It is natural to model the mean of *W*_*ik*_ as

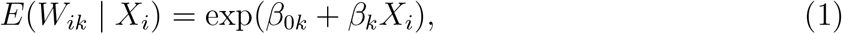

where *β*_0*k*_ is the intercept and *β*_*k*_ is the AA-level association. Sequencing generates counts via a (feature-specific biased) multinomial sampling mechanism, leading to

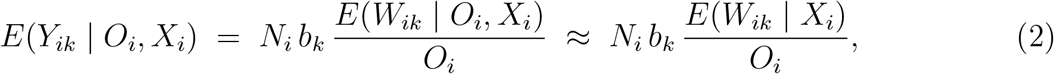

where *b*_*k*_ is a feature-level multiplicative factor accounting for measurement biases, such as differences in DNA extraction efficiency and primer binding/amplification efficiency in measuring various features^14^. In Additional file 1: Supplementary Methods 1.1, we show that the approximation *E*(*W*_*ik*_ | *O*_*i*_, *X*_*i*_) ≈ *E*(*W*_*ik*_ | *X*_*i*_) holds with typical microbiome data (large *K*, no overwhelmingly dominant feature, and no pervasive cross-feature dependence).

Since *O*_*i*_ is the summation of *W*_*ik*_’s, it will generally also be associated with *X*_*i*_ whenever any *W*_*ik*_ is associated with *X*_*i*_. Therefore, we re-parameterize *O*_*i*_ as 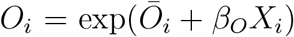, define the residual 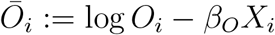, and express (2) as

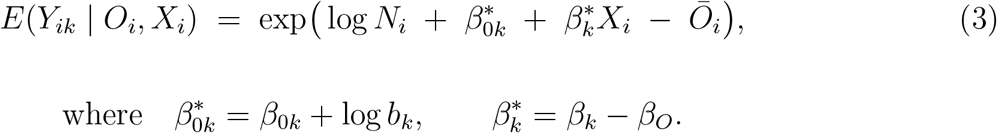

Since *O*_*i*_ is unobserved, we work with the marginal RA mean. We assume a mean-independence condition:

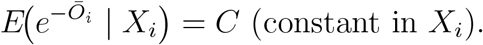

Then the marginal RA mean can be expressed as

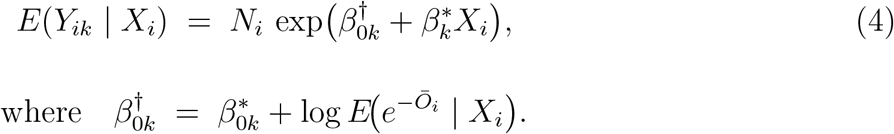

Thus, unidentifiable multiplicative factors (*b*_k_ and the load factor 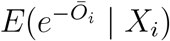) are absorbed into the intercept 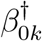, while the RA-level association 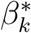 equals the AA-level association shifted by the common quantity *β*_*O*_ across all features.

#### Generating summary statistics for a study with independent samples

To estimate 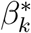, we propose using the quasi-Poisson regression with the RA mean specified in (4) and variance Var(*Y*_*ik*_) = *ϕE*(*Y*_*ik*_), where *ϕ* is the dispersion parameter. This semi-parametric approach accounts for the overdispersion of count data and does not rely on specific distributional assumptions. Let 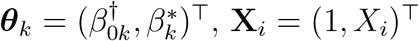, and 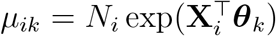. We employ the Firth bias-corrected^54^ Poisson quasi-score function

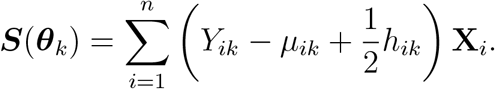

The **X**_*i*_ can be expanded to include potential confounders. The *h*_*ik*_ is the *i*th diagonal element of the ‘hat’ matrix ***H*** = ***W*** ^1/2^**X**(**X**^⊤^***W* X**)^−1^**X**^⊤^***W*** ^1/2^, where the ***W*** is a diagonal matrix with diagonal elements (*µ*_1*k*_, …, *µ*_*nk*_) and **X** is the design matrix. The Firth correction effectively reduces bias in parameter estimates, particularly in cases of small sample sizes and features with a high proportion of zero counts^54^.

We can estimate 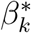 by solving ***S***(***θ***_*k*_) = **0**. However, fitting a separate model for each feature-covariate pair becomes computationally slow when the number of features and covariates is large. To address this, we propose estimating 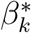 using the score statistic. Let 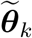 denote the estimate of ***θ***_*k*_ under the null model, which is independent of the covariate of interest. Partition the quasi-score function ***S***(***θ***_*k*_) into (*S*_1_, ***S***_0_), corresponding to the parameter of interest 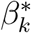 and the nuisance parameters, respectively. Similarly, partition the quasi-information matrix 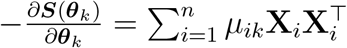as 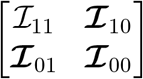. The RA-level association effect 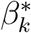 is estimated using the score statistic

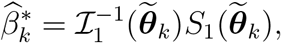

Where 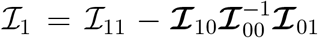. This estimate is derived from the Taylor series expansion of the quasi-score function. Score-statistic-based association analysis is widely used in modern GWAS due to its greater statistical accuracy and numerical stability compared to Wald-statistic-based analysis^55,56^.

The variance of 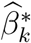 can be robustly estimated using the sandwich formula^57^

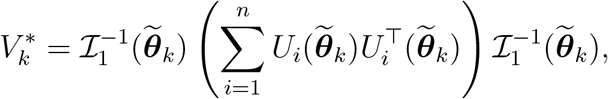

where 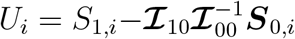 and ***S***_0,*i*_ are the *i*th summand of *S*_1_ and ***S***_0_, respectively.

#### Generating summary statistics for a study with correlated samples

For studies with correlated samples (e.g., longitudinal data, repeated measurements, or family-based data), the PALM summary statistics can be obtained in a similar manner, with adjusted variance estimates to account for correlation among samples within a cluster. Suppose we have *n* clusters in the data. Let *i* index clusters (*i* = 1, …, *n*) and *t* index correlated samples within a particular cluster (*t* = 1 …, *T*_*i*_). The notation for data is defined similarly to the independent sample case, but subscript *i* is replaced with *it* to index sample *t* within cluster *i*. The quasi-score function can be written as

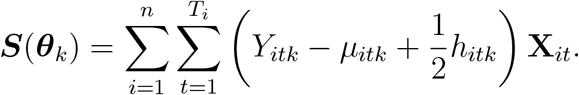

Let (*S*_1_, ***S***_0_) be the partition of ***S***(***θ***_*k*_) corresponding to the parameter of interest 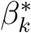 and the nuisance parameters. We estimate 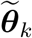, construct 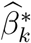, and calculate 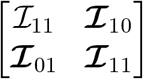 and ℐ_1_ using the same procedures described above. The sandwich estimator for the variance of 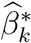 is adjusted to account for correlated samples as

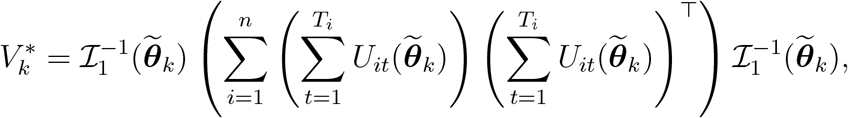

where 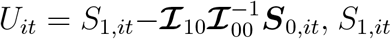 and ***S***_0,*it*_ are the *it*th summand of *S*_1_ and ***S***_0_, respectively.

#### Recovering AA-level summary statistics

We are interested in detecting AA-level associations (*H*_0_ : *β*_*k*_ = 0). Given the relationship between AA- and RA-level association effects, 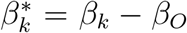, it follows that *β*_*O*_ represents the common compositional effect shared by all features (*k* = 1, …, *K*). Without additional assumptions, *β*_*O*_ is not estimable, and AA-level effects are unidentifiable from RA-level effects. Here, we leverage the high dimensionality of microbial features and assume that the majority of features have no AA-level effects (i.e., *β*_*k*_ = 0 for most *k*). Under this sparse AA signal assumption, we propose estimating −*β*_*O*_ using median 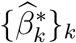 and obtaining the estimate of the AA-level effect as

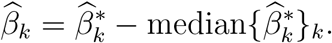

A key advantage of using the median is its robustness to extreme values and outliers.

The variance of 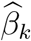 can be calibrated from 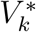 by taking into account the variability of the median as (Additional file 1: Supplementary Methods 1.2)^58^

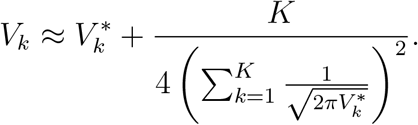

The 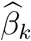 and *V*_*k*_ can be conveniently shared as AA-level summary statistics across studies and subsequently combined to perform meta-analysis for association testing.

#### Meta-analysis

Suppose we want to meta-analyze *L* microbiome association studies. For study *ℓ* = 1, …, *L*, we obtain AA-level association effect estimate 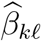 and its variance *V*_*kℓ*_ for each feature *k*. A feature may be unobserved or excluded from a given study due to its low prevalence in that study. We define ℒ_*k*_ as the set of indices for studies where feature *k* is observed. Under the fixed-effect model, the inverse-variance weighted estimate of the overall effect is

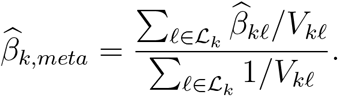

The variance of 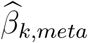 is estimated as

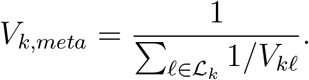

Finally, we construct the test statistic 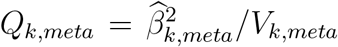 and calculate the *p*-value using the chi-square distribution with one degree of freedom.

### Comparison of methods

We compared the performance of PALM with ANCOM-BC2, DESeq2, LinDA, LM-CLR, and MaAsLin2. In all meta-analyses, both in simulations and real data applications, summary statistics for each method were generated at the individual study level. A fixed-effect meta-analysis was then performed by combining these summary statistics for association testing. In the simulation studies and the first two real data applications, microbial features associated with the covariate of interest were identified using the BH procedure with a target FDR of 0.05. In the third real data application (mbGWAS), we followed the mbGWAS convention^6,59,60^ and reported genome-wide significance at *p*-value *<* 5 *×* 10^−8^. All analyses were conducted in R (v4.3.1). Feature prevalence filters were applied externally, and no additional prevalence filters were applied within each method’s function.

- ANCOM-BC2: we used the `ancombc` function in the package ANCOMBC (v2.2.2) with the specified argument `prv_cut = 0`. Summary statistics `lfc_labels1` and `se_labels1` were extracted from the output.
- DESeq2: we used the R package DESeq2 (v1.48.2) with default settings, except that the estimation of size factors was set to use ‘`poscounts`’, following^61^. Differential abundance test was performed with the function `DESeq`, which uses the `nbinomWald` test function as the default. Summary statistics `log2FoldChange` and `lfcSE` were extracted from the output.
- LinDA: we used the `linda` function in the package MicrobiomeStat (v1.2) with the specified arguments `feature.dat.type =` ‘`count`’. Summary statistics `log2FoldChange` and `lfcSE` were extracted from the output.
- LM-CLR: we first added a pseudo-count (default: 0.5) to each cell of the count matrix and then applied the centered log-ratio transformation. To analyze a study with independent samples, we used the `lm` function available through the base R distribution for each transformed feature variable as the response. To analyze a study with correlated samples, we used the `lmer` function in the package lme4 (v1.1-34). Summary statistics were then extracted from the coefficient table of the output.
- MaAsLin2: we used the `Maaslin2` function in the package Maaslin2 (v1.14.1) with the specified arguments `min prevalence = 0`, `transform =` ‘`AST`’. Summary statistics `coef` and `stderr` were extracted from the output.

For all methods, the fixed-effect meta-analysis was performed using the function `rma` in the package metafor (v4.2-0) with specified argument `method=`‘`EE`’. The Cochran’s *Q* heterogeneity test *p*-value was also extracted from the output.

### Simulation strategy

We simulated microbiome data by mimicking the structure of the real metagenomics data from five studies for CRC^8^ using the package MIDASim (v0.1.0)^26^. MIDASim is a fast simulator that reproduces the distributional and correlation structure of a user-supplied template microbiome data. It supports controlled shifts in sequencing depth and means of microbial features. We considered *K* = 401 species with a prevalence of at least 20% across the five studies in the real data and *K* = 92 genera derived by collapsing the species-level data. To learn the structure of species (or genus) counts in each study, we used the `MIDASim.setup` function in nonparametric mode. The function returned the mean proportion vector (***µ***) and additional parameters (e.g., correlation matrices) that define the structure of the template data. We simulated two equally sized groups of samples (*X* = 1 and *X* = 0) by introducing differential AA signals into the mean proportion vector between the groups. Specifically, we randomly selected a subset of features as “active” features. Half of these were chosen randomly from relatively abundant features, defined as having an average proportion of at least 10^−3^ in the original data, while the other half were randomly selected from the remaining features. The mean proportion vectors for the two groups were initialized as ***µ***_1_ = ***µ***_0_ = ***µ***. For active features with positive effects, we increased their proportions in ***µ***_1_, and for active features with negative effects, we increased their proportions in ***µ***_0_. The fold increase for each active feature was randomly sampled from Uniform(1, Δ). After adjusting for all active features, ***µ***_1_ and ***µ***_0_ were normalized so that their elements summed to 1, ensuring they represented valid proportions. Note that the proportions of all features were shifted after this step. However, the active features were considered true differential AA features, driven by changes in the underlying AA between the two groups, which in turn caused the shift in RA for all features. Finally, we consolidated all parameters, including ***µ***_1_ and ***µ***_0_, the specified sequencing depths, and the other parameters learned from the template data, using the `MIDASim.modify` function. We then generated samples for the two groups using the `MIDASim` function. This procedure was repeated for each study, resulting in realistic simulated datasets closely resembling the original data (Additional file 1: Fig. S3). The randomly selected active features and their associated fold changes during the spike-in procedure were consistent across studies, representing homogeneous association effects.

We considered a broad range of scenarios in our simulations: (1) Two settings for feature size: species-level (*K* = 401) or genus-level (*K* = 92) data; (2) Two settings for sample sizes over the five studies: large sample size (*n* = 100, 120, 140, 160, 180) or small sample size (*n* = 20, 40, 60, 80, 100); The factor Δ, which controls the size of the fold change in proportion, was set to 5 for the large sample size setting and 10 for the small sample size setting. (3) Two effect direction settings: balanced positive/negative effects (active features had equal probabilities of having positive/negative effects) or dominant positive effects (all active features had positive effects). (4) Two sequencing depth unevenness settings: even (sequencing depths of all samples were randomly drawn from the pool of sequencing depths in the real CRC data) or uneven (further doubled the sequencing depth for the samples in a randomly picked group); (5) A range of active feature proportions from 0.05 to 0.2. Combining all these factors resulted in 64 simulation settings, and we generated 100 simulated datasets for each setting.

We conducted another set of simulations to meta-analyze studies with correlated samples. First, we simulated independent samples as described above. Next, we assigned every two samples *Y*_*i*1_ and *Y*_*i*2_ from the same group to form cluster *i*. To introduce correlation, we simulated another independent sample 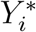 under the same group as *Y*_*i*1_ and *Y*_*i*2_. This independent sample was then mixed with *Y*_*i*1_ and *Y*_*i*2_ to create the correlated samples 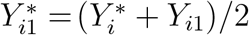 and 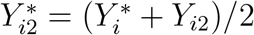 for cluster *i*.

### Analyzed datasets

#### Colorectal cancer (CRC) studies

The five studies include subjects from five different countries. The sample sizes of the studies are 109, 127, 120, 114, and 104 (Additional file 1: Table S1). All raw sequencing data across studies were reprocessed using the same bioinformatics pipeline for taxonomic profiling^8^. We removed two samples with an outlier sequencing depth of less than 2,000. Our analysis focused on *K* = 401 species after removing those with a prevalence of less than 20% (prevalence was defined as the percentage of non-zero counts in all samples across studies).

#### Metabolome (MTBL) studies

In all studies, the microbiome and metabolome were profiled from the same fecal samples. The datasets were curated and processed as part of an established effort^45^. We used the eight adult studies from this resource, with sample sizes of 96, 294, 240, 220, 444, 382, 287, and 164. These studies exhibit notable heterogeneity in study design, cohort demographics, and technical aspects of microbiome and metabolite measurements (Additional file 1: Table S3). Our analysis focused on 450 HMDB-annotated metabolites with study-specific prevalence ≥ 10% in at least two studies and 101 genera with study-specific prevalence ≥ 10% in at least one study. Following standard practice for normalizing skewed metabolite data, we applied a rank-based inverse normal transformation to the metabolite variables.

#### Host genetics study

The study is based on a population-based birth cohort of term-born and healthy infants^62^. The host genetics and microbiome data were collected from a subset of the cohort with 502 infants aged between 2 and 242 days. Most of the infants were non-Hispanic White. Genotypes were determined using the Illumina Genome Diversity Array (GDA, 8v1.0) at the Center for Inherited Disease Research (CIDR) (Baltimore, MD). Individuals were removed if they met any of the following quality control (QC) criteria: genotype vs. self-reported sex mismatches, unexpected duplicate samples, *>* 10% missing genotypes, and autosomal heterozygosity exceeding five standard deviations (SD=0.8%) above or below the mean of 30.4%. SNPs meeting any of the following QC were excluded: *>* 10% missing genotypes, exact Hardy-Weinberg *p*-value *<* 5 *×* 10^−8^, or genotype concordance *<* 90%. A total of 1,785,122 directly genotyped SNPs remained following QC. Ancestry principal components were estimated from autosomal SNPs with MAF *>* 0.005 and LD r2 *<* 0.10, using the 1,000 Genomes dataset as the reference. Imputation was performed using the TopMed reference panel (r2 version 1.0.0) of *>*100,000 whole genome sequences (human genome build 38) and the TopMed Imputation Server (version 1.7.3). Specifically, Eagle (version 2.4) was used for genotype phasing and minimac4 (version 1.0.2) was used for genotype imputation. Genotype dosages for 6,174,920 biallelic SNPs with MAF *>* 5% and imputation INFO score *>* 0.9 were included in the analysis.

The details of microbiome sample collection, processing, and sequencing were previously reported^63^ and were briefly summarized here. For the characterization of the gut microbiome, we extracted bacterial DNA from the stool samples collected at enrollment using the DNeasy PowerSoil Kit (Qiagen). The libraries were then sequenced on an Illumina MiSeq with 2*×*250 base pair reads. The 16S rRNA sequences were processed using the dada2 package, grouping the sequences into ASVs and assigning taxonomy based on the SILVA reference database. Our analysis focused on 109 ASVs with a prevalence greater than 5%.

## Declarations

### Funding

This work was supported by the NIH grants R01AI192972, R01GM140464, UG3/UH3 OD023282, U19 AI 095227, UG3 OD035516, UL1 TR002243, and NSF grant DMS-2054346.

### Availability of data and materials

The data for the CRC studies were obtained from https://github.com/zellerlab/crc_meta^64^. The data for the MTBL studies were obtained from https://github.com/borenstein-lab/microbiome-metabolome-curated-data^65^. Individual-level genotype data for the INSPIRE birth cohort analyzed in this study are available through the NIH database of Genotypes and Phenotypes (dbGaP) under controlled access, accession phs004036.v1.p1^66^. PALM is implemented in the R package PALM, publicly available at https://github.com/ZjpWei/PALM_package^67^. The site also contains a complete set of manuals and instructions. The scripts generating reported results are deposited at https://github.com/ZjpWei/PALM^68^ and Zenodo https://doi.org/10.5281/zenodo.17547702^69^ under the GPL-3.0 license.

### Authors’ contributions

ZZT conceived the idea and oversaw the study. ZZT developed PALM with contributions from GC and ZW. ZW implemented the method, conducted numerical studies, and developed the PALM R package. QH conducted the mbGWAS analysis. TVH, CR, SRD, MHS, and AML contributed to the data generation for the mbGWAS analysis. ZZT wrote the first version of the manuscript and all authors contributed to the writing. All authors read and approved the final manuscript.

## Acknowledgements

The authors thank the editors and three anonymous reviewers for their insightful comments and suggestions, which helped improve the quality of the manuscript.

## Peer review information

Andrew Cosgrove, Claudia Feng, and Zhenrun Jerry Zhang were the primary editors of this article and managed its editorial process and peer review in collaboration with the rest of the editorial team. The peer-review history is available in the online version of this article.

## Ethics approval and consent to participate

Not applicable

## Consent for publication

Not applicable

## Competing interests

Not applicable

## Supplementary Information

Additional file 1: Supplementary Methods 1.1-1.3, Figs S1-S14, and Tables S1, S3.

Additional file 2: Tables S2.

Additional file 3: Tables S4.

## Notes

### Competing Interest Statement

The authors have declared no competing interest.

