## Supplementary Material for "Fast and reliable association discovery in large-scale microbiome studies and meta-analyses using PALM"

### 1 Supplementary Methods

#### 1.1 Justification of $E(W_{ik} \mid O_i, X_i) \approx E(W_{ik} \mid X_i)$

The linear projection (best linear predictor) of $E(W_{ik} \mid O_i, X_i = x)$ on $O_i$ gives

$$E(W_{ik} \mid O_i, X_i = x) \;\approx\; \mu_k(x) + \lambda_k(x)\{O_i - \mu_O(x)\},$$

where $\mu_k(x) := E(W_{ik} \mid X_i = x)$, $\mu_O(x) := E(O_i \mid X_i = x)$, $v_O(x) := \mathrm{Var}(O_i \mid X_i = x)$, and

$$\lambda_k(x) := \frac{\mathrm{Cov}(W_{ik}, O_i \mid X_i = x)}{\mathrm{Var}(O_i \mid X_i = x)}.$$

Thus the deviation from $\mu_k(x)$ is controlled by $\lambda_k(x)$ and by the concentration of $O_i$ around $\mu_O(x)$. All constants below do not depend on $K$, $k$, or $x$.

**Assumptions:**

**(M1) No dominant feature / linear mean growth.** There exist $0 < c_\mu \le C_\mu < \infty$ such that

$$\frac{1}{K}\sum_{k=1}^{K} \mu_k(x) \in [c_\mu, C_\mu].$$

Equivalently, $\mu_O(x) = \sum_{k=1}^{K} \mu_k(x)$ grows linearly in $K$ and $\max_k \mu_k(x)/\mu_O(x) \le C_*/(c_\mu K)$ for some $C_* < \infty$.

**(M2) Bounded marginal variance and no pervasive cross-feature dependence.** Let $\Sigma_{kj}(x) := \mathrm{Cov}(W_{ik}, W_{ij} \mid X_i = x)$ and $R_k(x) := \sum_{j \ne k} |\Sigma_{kj}(x)|$. There exist $0 < C_v, C_c < \infty$ such that

$$\max_k \mathrm{Var}(W_{ik} \mid X_i = x) \le C_v, \qquad \max_k R_k(x) \le C_c.$$

This allows block/local correlation but rules out uniformly strong dense dependence.

**(M3) Concentration of total load (consequence of (M1)–(M2)).** Under (M1)–(M2), the total load variance grows at most linearly: $v_O(x) = \mathcal{O}(K)$, while $\mu_O(x)$ grows linearly with $K$. Hence the coefficient of variation $\mathrm{CV}^2(O_i \mid X_i = x) = v_O(x)/\mu_O(x)^2 = \mathcal{O}(1/K)$.

Under (M2), $\mathrm{Cov}(W_{ik}, O_i \mid X_i = x) = \mathrm{Var}(W_{ik} \mid X_i = x) + \sum_{j \neq k} \Sigma_{kj}(x) = \mathcal{O}(1)$, and, by (M1)–(M2), $v_O(x) = \mathrm{Var}(O_i \mid X_i = x) = \mathcal{O}(K)$ because the variance of the sum adds at most linearly under bounded row sums and bounded marginal variances. Therefore,

$$\lambda_k(x) = \frac{\mathrm{Cov}(W_{ik}, O_i \mid X_i = x)}{\mathrm{Var}(O_i \mid X_i = x)} = \mathcal{O}\left(\frac{1}{K}\right).$$

By Chebyshev's inequality and (M3), for any $c > 0$,

$$\Pr\Big(|O_i - \mu_O(x)| \leq c\sqrt{v_O(x)} \;\Big|\; X_i = x\Big) \;\geq\; 1 - \frac{1}{c^2}.$$

On this event,

$$\big|E(W_{ik} \mid O_i, X_i = x) - \mu_k(x)\big| \;\leq\; |\lambda_k(x)|\,|O_i - \mu_O(x)| \;\leq\; \frac{C_\lambda}{K} c\sqrt{v_O(x)} \;=\; \mathcal{O}\left(\frac{1}{\sqrt{K}}\right).$$

Hence, for any $\varepsilon > 0$, there exists a finite number $C_\varepsilon > 0$ (depending on $\varepsilon$ but not on $K$, $k$, or $x$) such that

$$\Pr\Big(\big|E(W_{ik} \mid O_i, X_i = x) - \mu_k(x)\big| \;\leq\; C_\varepsilon K^{-1/2} \;\Big|\; X_i = x\Big) \;\geq\; 1 - \varepsilon.$$

Thus, the difference between $E(W_{ik} \mid O_i, X_i = x)$ and $E(W_{ik} \mid X_i = x)$ is of order $K^{-1/2}$ in probability as $K$ increases.

#### 1.2 Derivation of the variance of $\widehat{\beta}_k$

Let $V_k$ and $V_k^*$ denote the variance of $\widehat{\beta}_k$ and $\widehat{\beta}_k^*$, respectively. Since $\widehat{\beta}_k = \widehat{\beta}_k^* - \text{median}\{\widehat{\beta}_k^*\}_k$, we have $V_k \approx V_k^* + \text{Var}(\text{median}\{\widehat{\beta}_k^*\}_k)$, provided that the dependence among $\widehat{\beta}_k^*$ is weak. We derive the formula for $\text{Var}(\text{median}\{\widehat{\beta}_k^*\}_k)$ as follows.

From quasi-likelihood theory[1], the estimate $\widehat{\beta}_k^*$ approximately follows a normal distribution with mean $\beta_k^* = \beta_k - \beta_O$ and variance estimated by $V_k^*$. For large $K$, the sample median is asymptotically normally distributed[2]

$$\text{median}\{\widehat{\beta}_k^*\}_k \sim \mathcal{N}(\beta^*_{\text{median}}, \text{Var}(\text{median}\{\widehat{\beta}_k^*\}_k)),$$

where $\beta^*_{\text{median}} = \text{median}\{\beta_k^*\}_k$. For normally distributed $\widehat{\beta}_k^*$'s, the $\text{Var}(\text{median}\{\widehat{\beta}_k^*\}_k)$ can be approximated as

$$\text{Var}(\text{median}\{\widehat{\beta}_k^*\}_k) \approx \frac{1}{4K\left(\frac{1}{K}\sum_{k=1}^{K} f_k(\beta^*_{\text{median}})\right)^2},$$

where $f_k(\beta^*_{\text{median}})$ is the normal density function of $\widehat{\beta}_k^*$ evaluated at $\beta^*_{\text{median}}$. By substituting the density function and simplifying the formula, we have

$$\text{Var}(\text{median}\{\widehat{\beta}_k^*\}_k) \approx \frac{K}{4\left(\sum_{k=1}^{K} \frac{1}{\sqrt{2\pi V_k^*}} \exp(-\frac{(\beta^*_{\text{median}}-\beta_k^*)^2}{2V_k^*})\right)^2}.$$

Under the sparse signal assumption that the majority of the features have no AA-level effects (i.e., $\beta_k = 0$ for most $k$), the majority of $\beta_k^*$ shares the same mean $-\beta_O$, leading to $\beta^*_{\text{median}} \approx -\beta_O$. Therefore, the exponential term approaches 1 for most $k$, resulting in the final variance approximation

$$\text{Var}(\text{median}\{\widehat{\beta}_k^*\}_k) \approx \frac{K}{4\left(\sum_{k=1}^{K} \frac{1}{\sqrt{2\pi V_k^*}}\right)^2}.$$

#### 1.3 An alternative method for estimating compositional effect

Suppose we want to meta-analyze $L$ independent microbiome association studies. Let $\boldsymbol{\beta}_{\bullet\ell} = \{\beta_{k\ell}\}$ and $\boldsymbol{\beta}^*_{\bullet\ell} = \{\beta^*_{k\ell}\}$ respectively denote the AA- and RA-level association effects for study $\ell$. The length of these coefficients could differ among studies because a feature may not be observed in all studies. Let $\mathcal{T}_\ell$ be the index set of features observed in study $\ell$ with size $K_\ell$. For each study $\ell$, we obtain RA-level association effect estimates denoted by $\widehat{\boldsymbol{\beta}}^*_{\bullet\ell} = \{\widehat{\beta}^*_{k\ell}\}_{k\in\mathcal{T}_\ell}$ and the diagonal matrix $\boldsymbol{V}^*_\ell$ in which the diagonal elements are the variance of $\widehat{\boldsymbol{\beta}}^*_{\bullet\ell}$. Let $\mathcal{T}_0$ denote the union set of all $\mathcal{T}_\ell$'s, $K_0$ denote the size of $\mathcal{T}_0$, and $\boldsymbol{\mu} = \{\mu_k\}_{k\in\mathcal{T}_0}$ be the collection of the meta AA-level association effects for all the features appear in at least one study. We introduce a hyperparameter $\delta_\ell$ to correct the compositional effect in $\widehat{\boldsymbol{\beta}}^*_{\bullet\ell}$ and propose to estimate sparse $\boldsymbol{\mu}$ by minimizing the best-subset-selection objective function

$$\mathcal{L}(\boldsymbol{\mu}) = n_0^{-1}\sum_{\ell=1}^{L}(\widehat{\boldsymbol{\beta}}^*_{\bullet\ell} + \delta_\ell\mathbf{1}_\ell - \boldsymbol{\mu}_{\mathcal{T}_\ell})^{\mathrm{T}}\boldsymbol{V}_\ell^{*-1}(\widehat{\boldsymbol{\beta}}^*_{\bullet\ell} + \delta_\ell\mathbf{1}_\ell - \boldsymbol{\mu}_{\mathcal{T}_\ell}), \text{ subject to } \|\boldsymbol{\mu}\|_0 \le s, \tag{1}$$

where $n_0$ is the total sample size across studies, $\mathbf{1}_\ell$ is a $K_\ell$-vector of ones, $\boldsymbol{\mu}_{\mathcal{T}_\ell}$ is the subset of $\boldsymbol{\mu}$ for features in $\mathcal{T}_\ell$, and $\|\boldsymbol{\mu}\|_0 = \sum_{k=1}^{K_0} I(\mu_k \neq 0)$ is the $\ell_0$ norm of $\boldsymbol{\mu}$. The unidentifiability of $\delta_\ell$ leads to infinite possible association profiles and we assume that true AA-level association profile involves the most parsimonious sets of features. Given $\delta_1, \ldots, \delta_L$ and $s$, we solve the best subset selection problem (1). The optimal values of the hyperparameters are determined by minimizing BIC defined as

$$\mathrm{BIC}(s, \delta_1, \ldots, \delta_L) = n_0^{-1}\sum_{\ell=1}^{L}(\widehat{\boldsymbol{\beta}}^*_{\bullet\ell} + \delta_\ell\mathbf{1}_\ell - \widetilde{\boldsymbol{\mu}}_{\mathcal{T}_\ell})^{\mathrm{T}}\boldsymbol{V}_\ell^{*-1}(\widehat{\boldsymbol{\beta}}^*_{\bullet\ell} + \delta_\ell\mathbf{1}_\ell - \widetilde{\boldsymbol{\mu}}_{\mathcal{T}_\ell}) + \|\widetilde{\boldsymbol{\mu}}\|_0\frac{\log n_0}{n_0}, \tag{2}$$

where $\widetilde{\boldsymbol{\mu}}$ is the estimate of $\boldsymbol{\mu}$ under a specification of hyperparameters. Details on solving the best subset selection problem and hyperparameter tuning algorithms were provided in[3]. Given the optimal hyperparameters $\widehat{\delta}_1, \ldots, \widehat{\delta}_L$, we obtain the estimate of the AA-level effect in study $\ell$ as $\widehat{\beta}_{k\ell} = \widehat{\beta}^*_{k\ell} + \widehat{\delta}_\ell$, $k = 1, \ldots, K$.

#### 2. Supplementary Figures and Tables

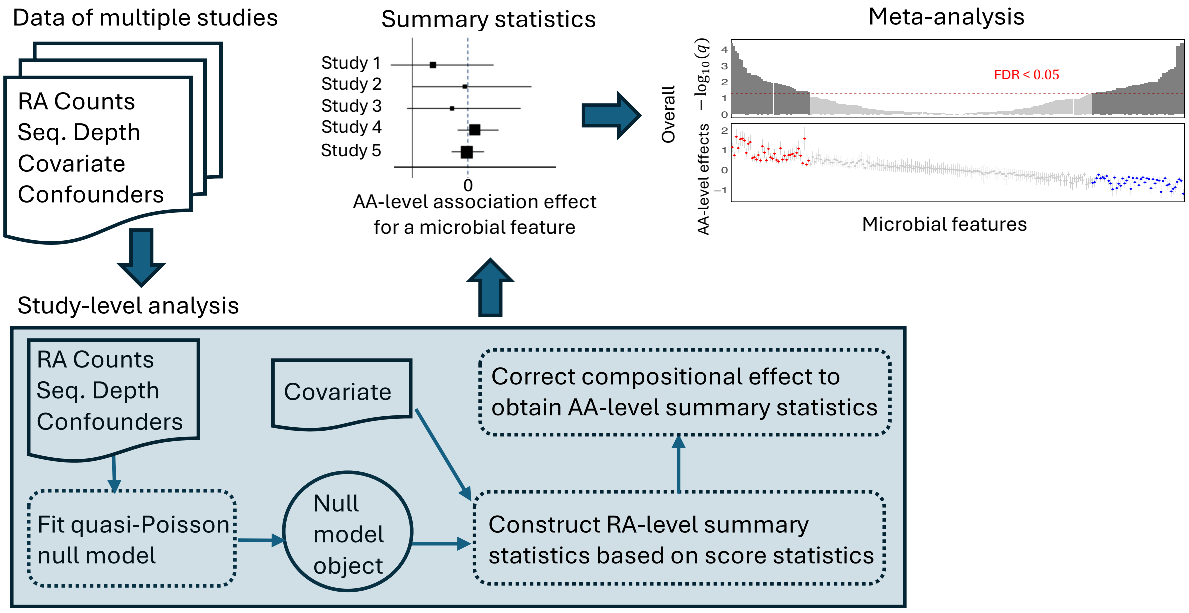


**Fig. S1: Schematic overview of PALM for meta-analysis.** For each study, PALM takes count data of microbial features, sequencing depth, the covariate of interest, and any relevant confounders as input to generate AA-level summary statistics. Specifically, for each feature, PALM first fits a quasi-Poisson null model (without the covariate of interest), then calculates score statistics based on the null model output and the covariate, and finally constructs the RA-level summary statistics (i.e., RA association effect estimate and variance) based on the score statistics. Next, PALM corrects the compositional effect using the RA-level summary statistics of all features to derive AA-level summary statistics. The forest plot illustrates the AA-level summary statistics for a feature. These summary statistics can be combined via meta-analysis to obtain the overall effect estimate, variance, and $p$-value. Covariate-associated microbial features are selected based on the test $p$-values, accounting for multiple testing across features.

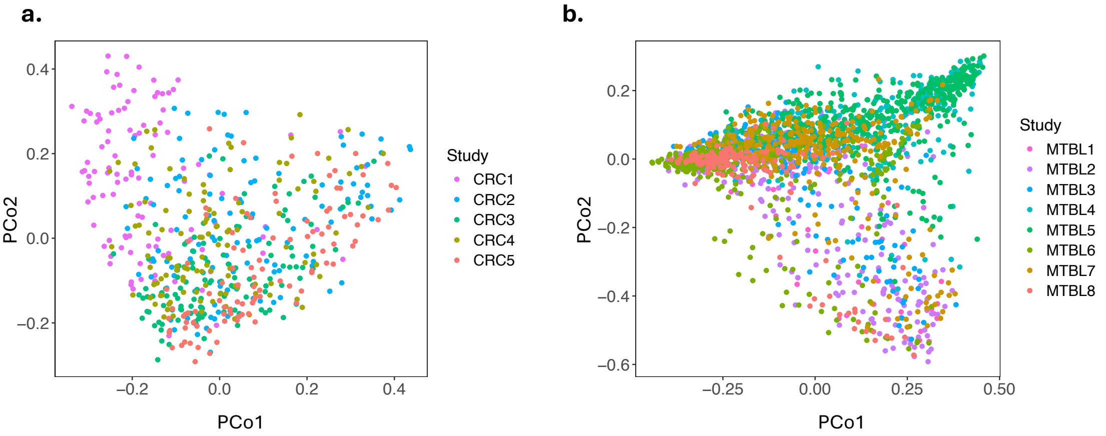


**Fig. S2: Principal coordinate plots of real microbiome data over studies in the two meta-analysis applications.** **a**, Meta-analysis of five microbiome-colorectal cancer (CRC) association studies. **b**, Meta-analysis of eight microbiome-metabolome (MTBL) association studies. Bray-Curtis dissimilarity was used in the principal coordinate analysis. Data points from different studies are color-coded. The PERMANOVA test $p$-values for testing the study effect in panels a and b are less than $10^{-5}$.

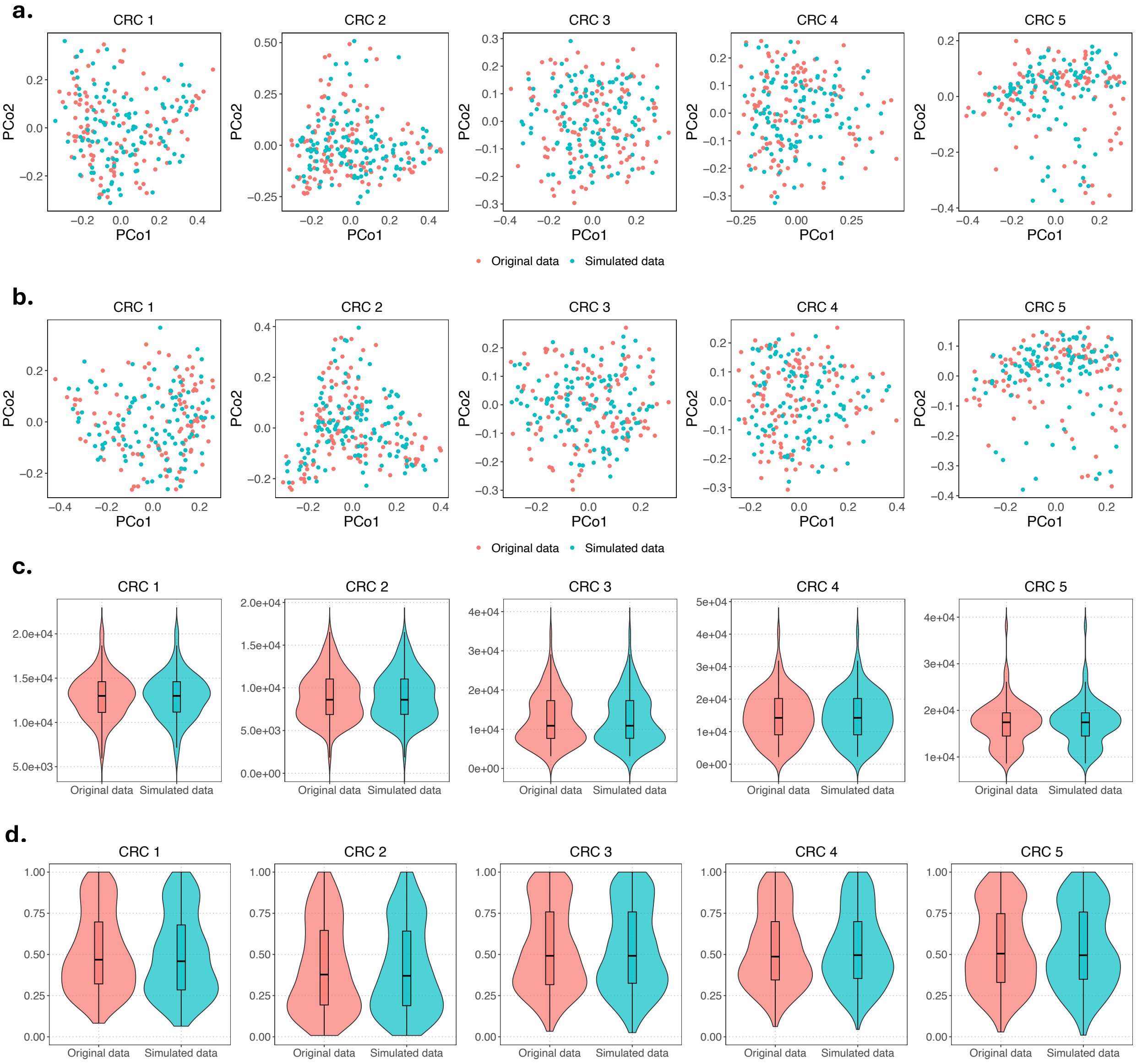


**Fig. S3: Comparison of the original and the simulated microbiome data.** Each panel represents a single study. The original data refers to the real data in a colorectal cancer (CRC) study (indicated in the panel title), and the simulated data was generated to mimic the original data using the MIDASim simulator. **a**, Principal coordinate plots based on Bray-Curtis dissimilarity for the original and simulated microbiome data. **b**, Principal coordinate plots based on Jaccard dissimilarity for the original and simulated microbiome data. The PERMANOVA test $p$-values for testing the difference between the original and simulated data exceed 0.1 for all principal coordinate plots, indicating a close resemblance between the two datasets. **c**, Sequencing depth across samples in the original and simulated microbiome data. **d**, Sample prevalence across features in the original and simulated microbiome data. The boxplots show the median (center line), the first and third quartiles (box edges), and whiskers extending to 1.5 times the interquartile range. Violin plots illustrate the density distribution of the data points.

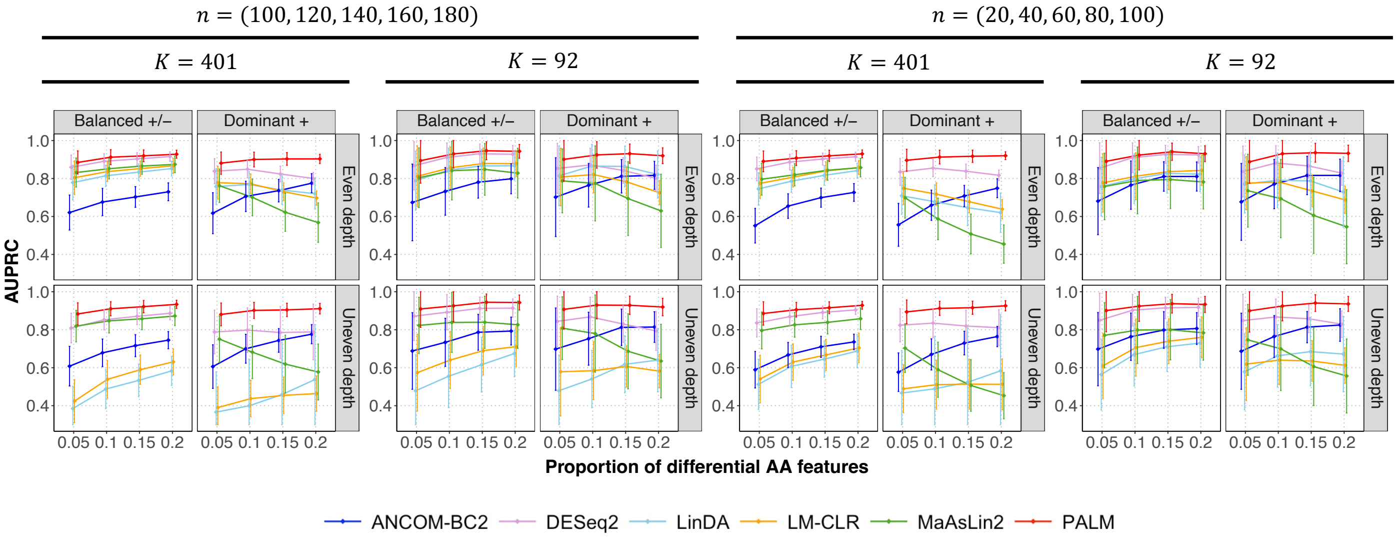


**Fig. S4: AUPRC comparison of different methods in the meta-analysis of five simulated studies with independent samples.** The x-axis represents the proportion of differential absolute abundance (AA) features. The top title shows the two scenarios of sample sizes $n$ in five studies. The second title shows the two scenarios of feature size $K$. The column facet title represents two effect direction scenarios: Balanced $+/-$, where differential AA features have an equal probability of exhibiting positive or negative effects, and Dominant $+$, where all differential AA features exhibit positive effects. The row facet title represents two sequencing depth scenarios: one with even sequencing depth and one with uneven sequencing depth between the comparison groups. Each panel displays the mean estimated area under the precision–recall curve (AUPRC) with $\pm$ standard errors (indicated by error bars) based on 100 simulation replicates.

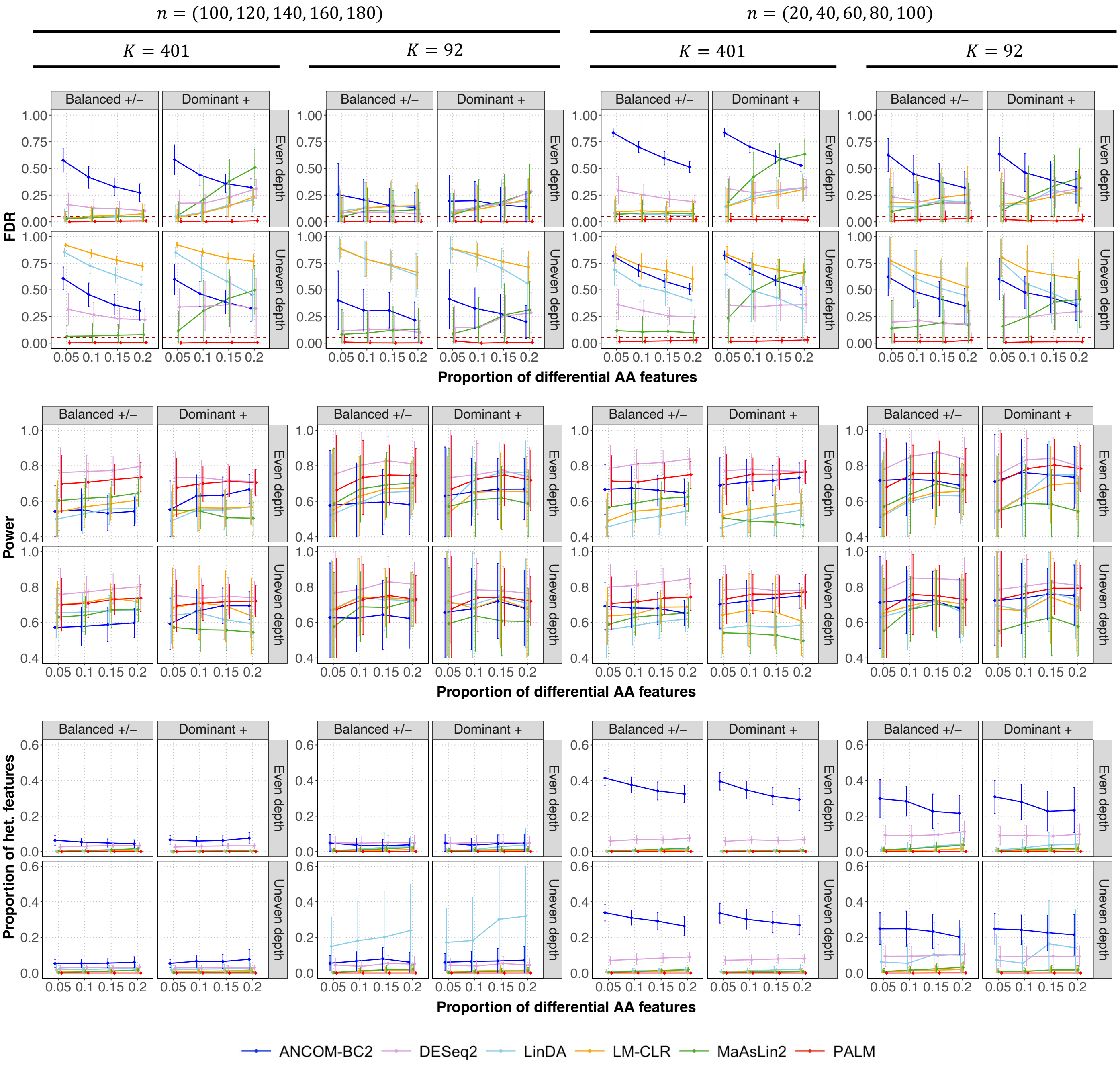


**Fig. S5: Comparison of methods in a meta-analysis of five independent simulated studies, with evaluation focused on rare features.** This figure has the same format as Fig. 1 but focuses on features with an average proportion below $10^{-3}$ in the template dataset.

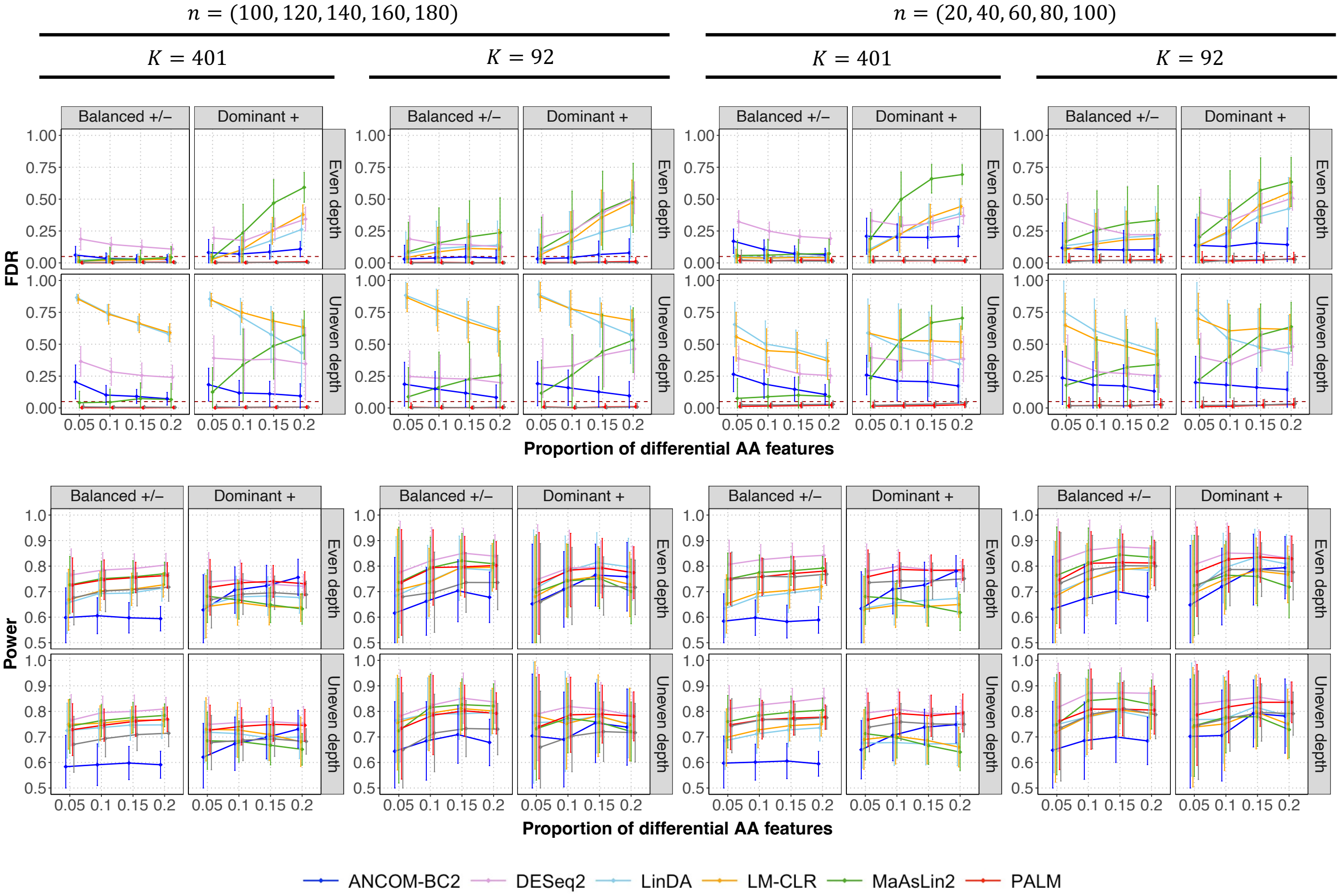


**Fig. S6: Comparison of PALM meta-analysis with pooled-data analysis by other methods across five simulated studies with independent samples.** The two sections of the figure show empirical FDR (top, the horizontal line indicates the target FDR) and power (bottom). The x-axis represents the proportion of differential absolute abundance (AA) features. The top title shows the two scenarios of sample sizes $n$ in five studies. The second title shows the two scenarios of feature size $K$. The column facet title represents two effect direction scenarios: Balanced $+/-$, where differential AA features have an equal probability of exhibiting positive or negative effects, and Dominant $+$, where all differential AA features exhibit positive effects. The row facet title represents two sequencing depth scenarios: one with even sequencing depth and one with uneven sequencing depth between the comparison groups. Each panel displays the mean estimated metrics with $\pm$ standard errors (indicated by error bars) based on 100 simulation replicates.

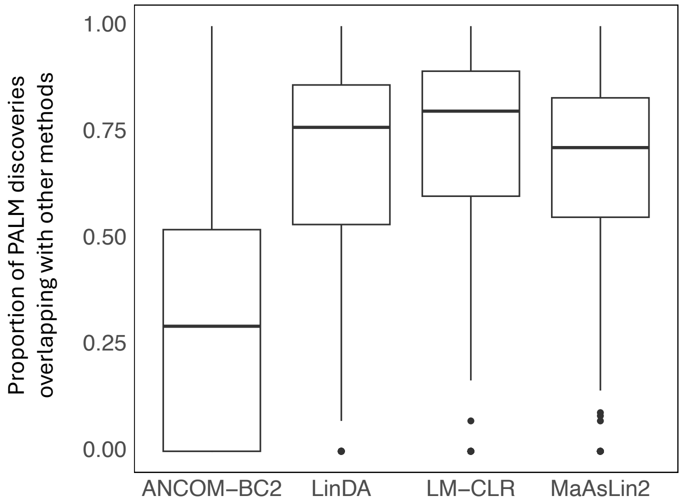


**Fig. S7: Overlap between PALM discoveries and other methods' discoveries in the meta-analysis of microbiome-metabolome association studies.** The x-axis lists the individual methods being compared. The y-axis shows the proportion of PALM-identified metabolite-associated features that overlap with those identified by other methods across all metabolites. Each boxplot indicates the median (center line), the first and third quartiles (box edges), whiskers extending to 1.5 times the interquartile range, and individual outliers (dots).

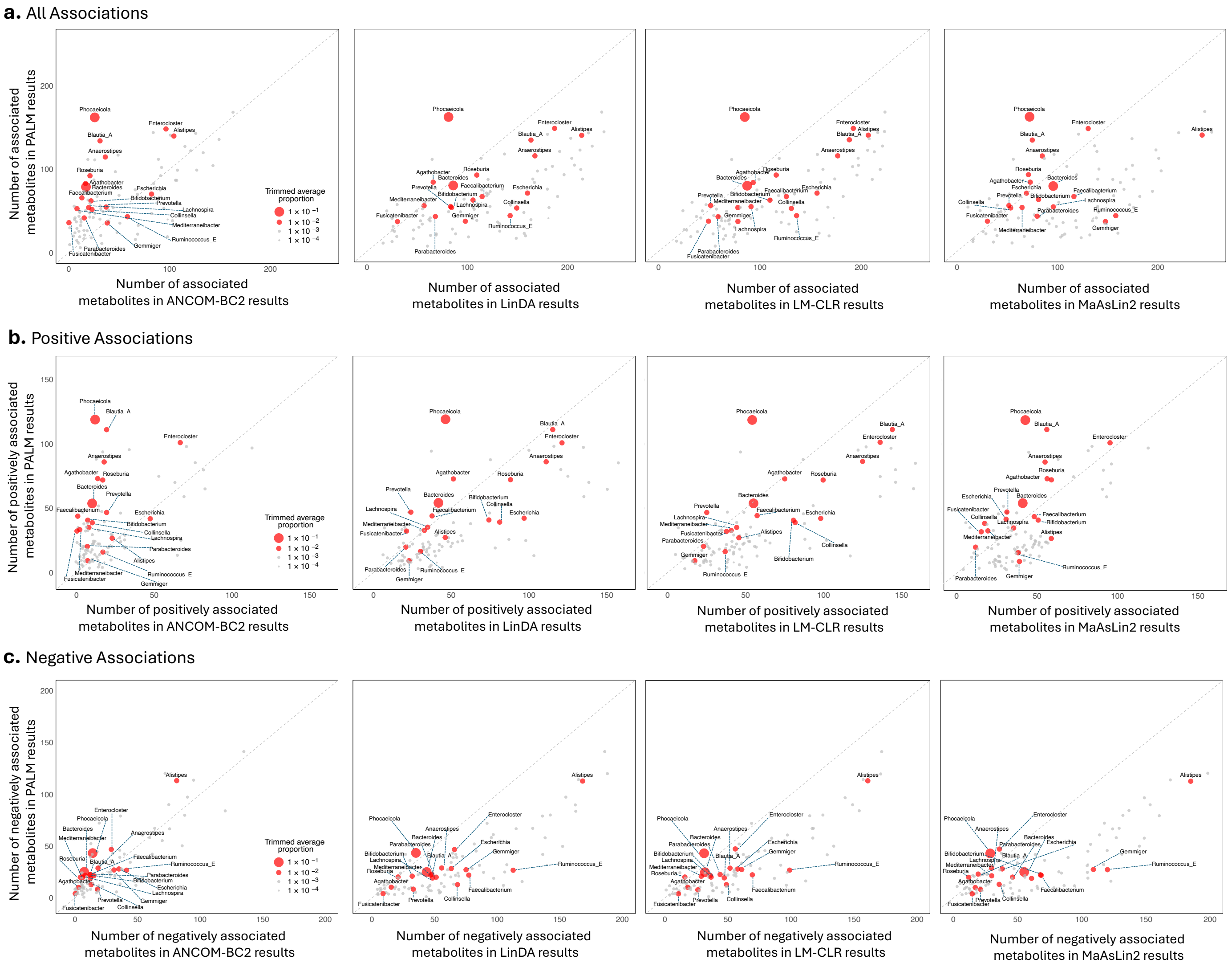


**Fig. S8: Number of associated metabolites per microbial feature in PALM results compared to other methods' results from the meta-analysis of microbiome-metabolome association studies.** **a**, number of all associations. **b**, number of positive associations. **c**, number of negative associations. Each dot represents a feature, with dot size proportional to the trimmed average proportion of the feature across all individuals in all studies. Features with an average proportion ≥ 0.01 are shown in red and annotated with their feature names, while other features are shown in gray. Compared to other methods, the top features in PALM results tend to have higher average proportions across all individuals in all studies (i.e., in **a**, the features that are well below diagonal lines have low proportions, while features closer to or above the diagonal line have higher proportions). This trend is driven by positive associations, rather than negative associations (see **b, c**).

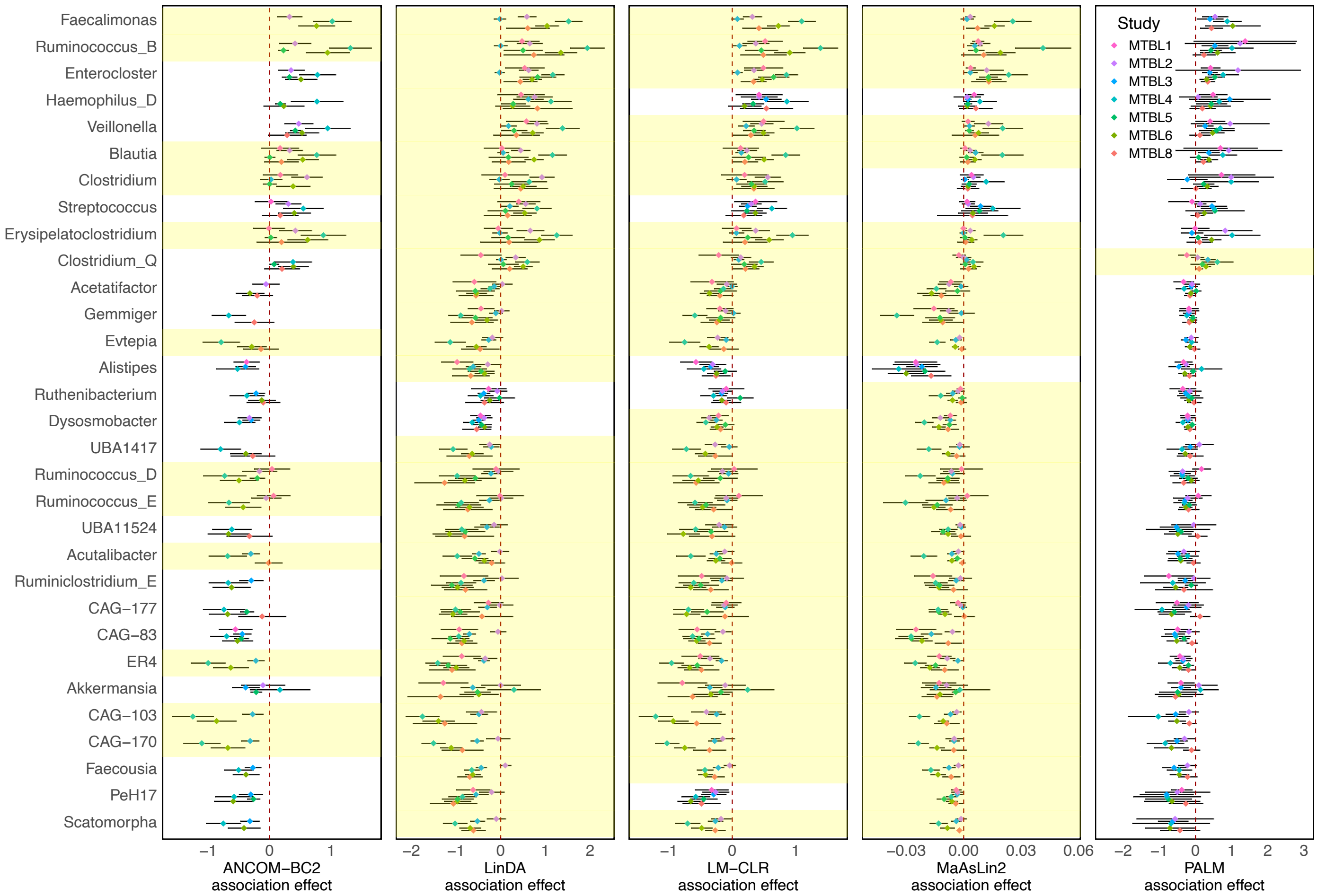


**Fig. S9: Forest plots for a set of microbial features associated with the metabolite cholic acid in the meta-analysis of microbiome-metabolome (MTBL) association studies.** x-axis represents a core set of cholic-acid-associated features consistently identified by all methods but exhibiting significant heterogeneity in effects across studies in at least one method. Each panel shows the summary statistics of a method, including the association effect estimates across studies (color-coded dots) and their 95% confidence intervals (black lines). The row for features with significant heterogeneous effects across studies is shaded yellow in each panel (Cochran's $Q$ test $q$-value $< 0.1$);

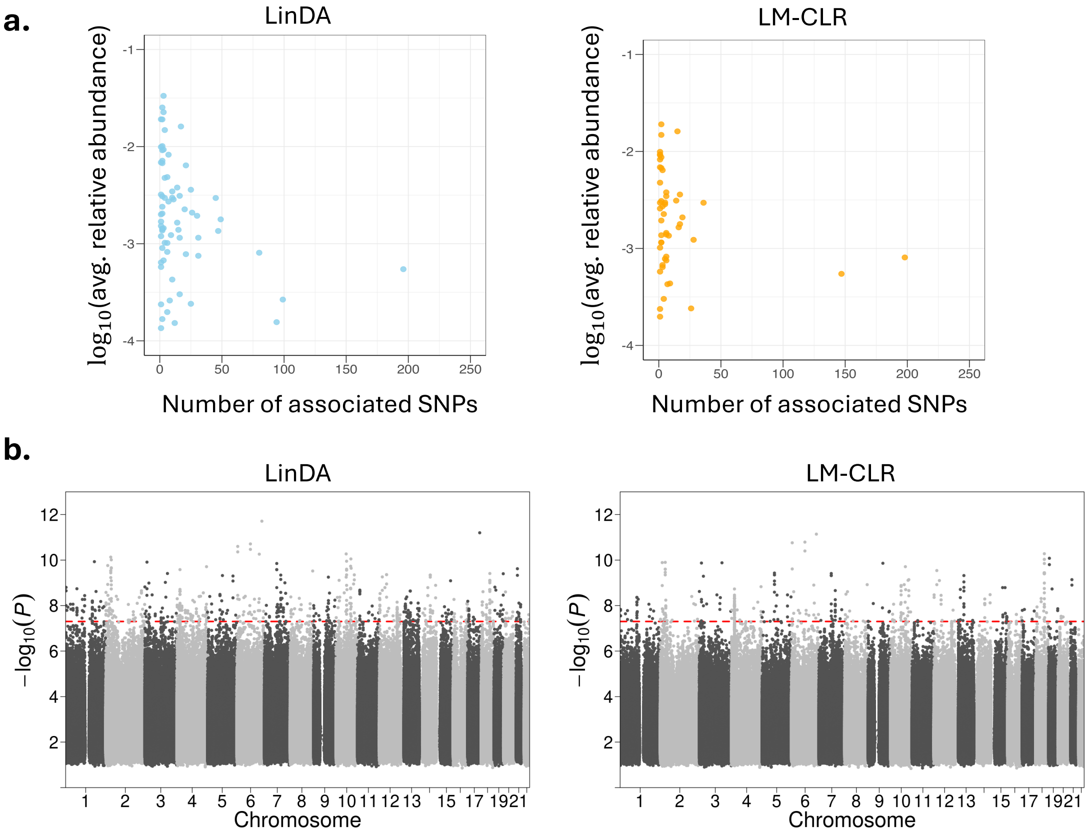


**Fig. S10: Additional microbiome and host genetics GWAS (mbGWAS) results from LinDA and LM-CLR.** **a**, Summary of the LinDA and LM-CLR identified SNP-associated ASVs (dots) shown in Fig. 6. The x-axis shows the number of SNPs associated with each ASV, and the y-axis shows the $-\log_{10}$ average relative abundance of each ASV. **b**, mbGWAS Manhattan plots for LinDA and LM-CLR when the pseudo-count of 0.01 was used in the analysis. Each SNP was tested against each of the 109 ASVs and the Manhattan plot shows the lowest resulting $p$-value for each SNP. The horizontal dashed line marks the genome-wide significance ($p$-value $= 5 \times 10^{-8}$).

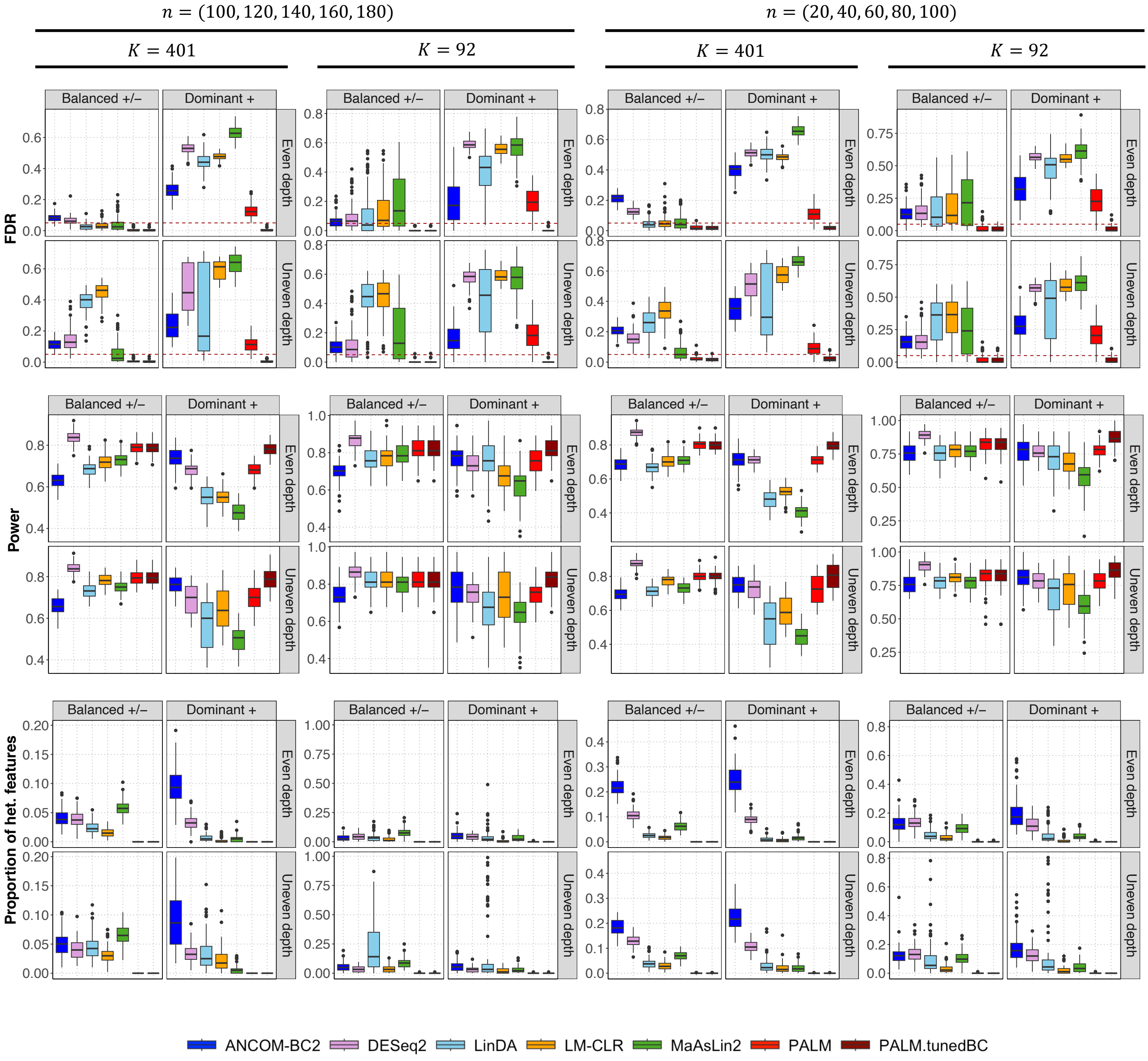


**Fig. S11: Comparison of different compositional effect correction approaches under the PALM model in a meta-analysis of five simulated studies with independent samples and dense differential AA features.** This figure has the same simulation setup and format as Fig. 1, except that the proportion of differential features is 40%. PALM uses median to correct compositional bias and PALM.tunedBC uses an alternative approach described in Supplementary Methods 1.3.

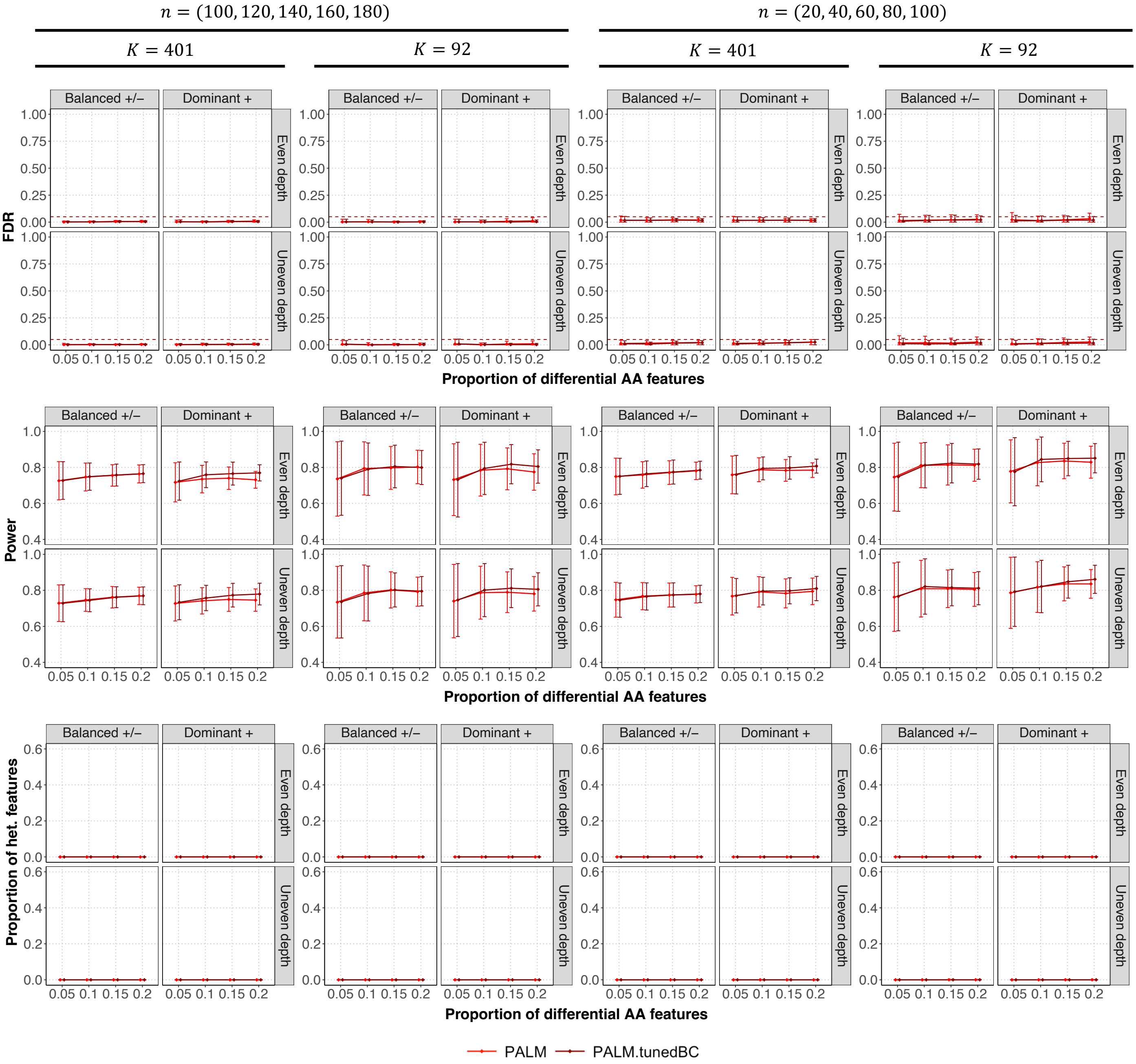


**Fig. S12: Comparison of different compositional effect correction approaches under the PALM model in a meta-analysis of five simulated studies with independent samples.** This figure has the same simulation setup and format as Fig. 1. PALM uses median to correct compositional bias and PALM.tunedBC uses an alternative approach described in Supplementary Methods 1.3.

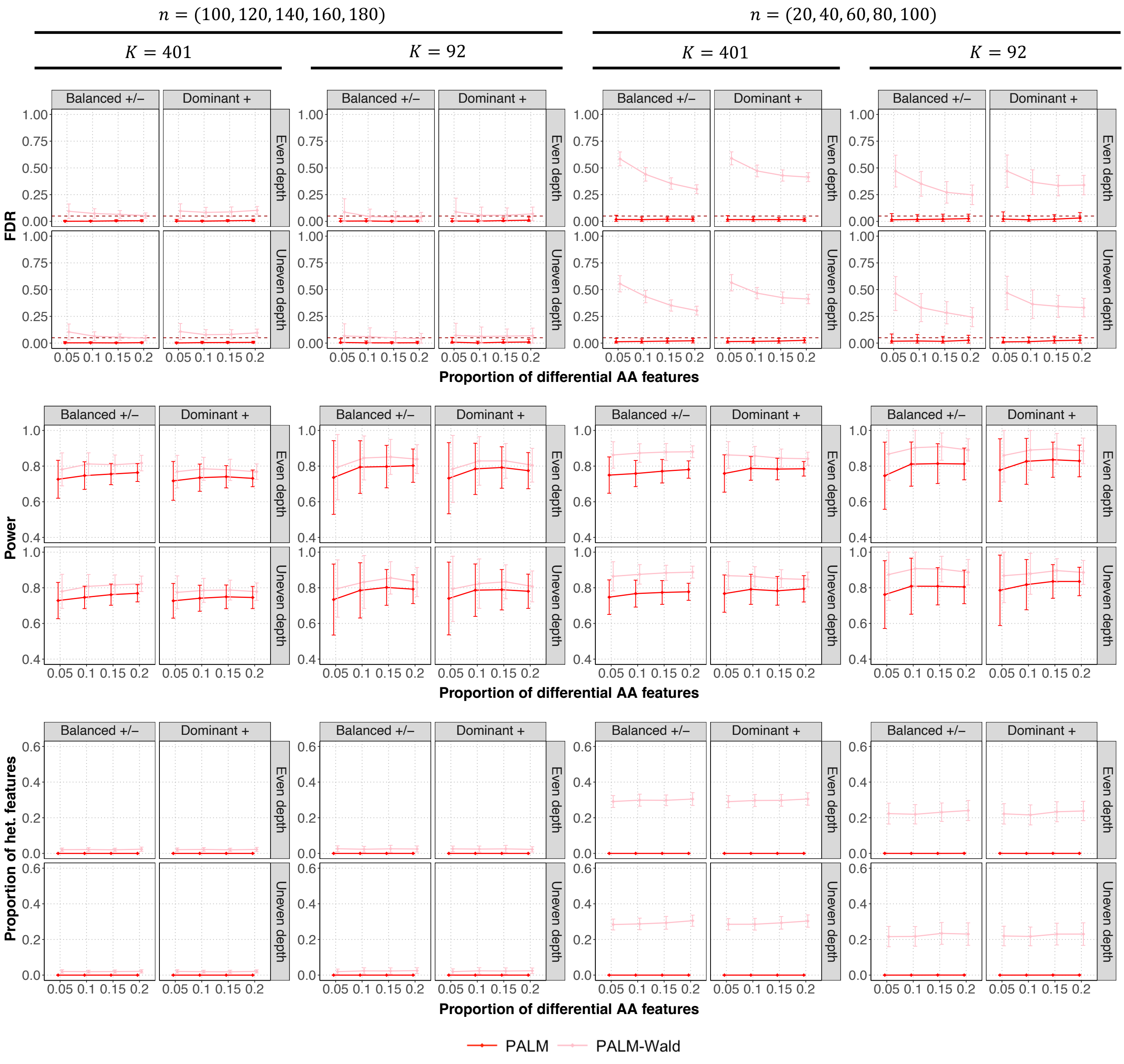


**Fig. S13: Comparison of the score-statistic-based test and the Wald test under the PALM model in a meta-analysis of five simulated studies with independent samples.** Similar to the score-statistic-based test, the Wald test also uses the robust sandwich estimator for variance estimation and applies the median-based compositional bias correction. This figure has the same simulation setup and format as Fig. 1.

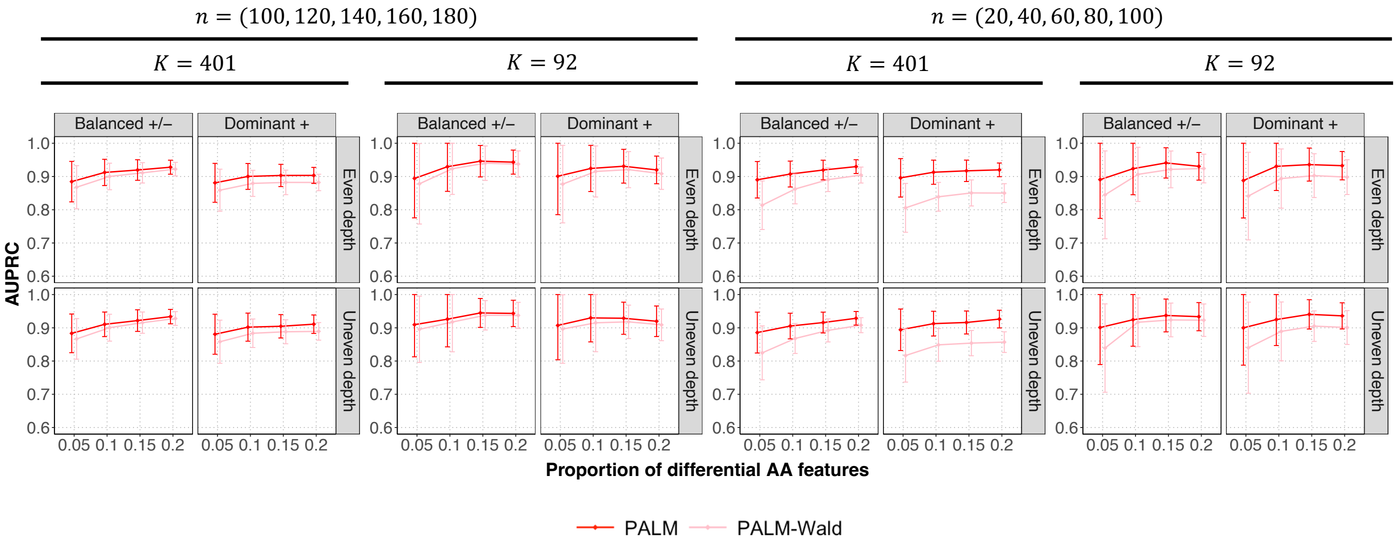


**Fig. S14: AUPRC comparison between the score-statistic–based test and the Wald test under the PALM model in a meta-analysis of five simulated studies with independent samples.** Each panel displays the mean estimated area under the precision–recall curve (AUPRC) with ± standard errors (indicated by error bars) based on 100 simulation replicates.

**Table S1**: Datasets of microbiome association studies for colorectal cancer.

| Study ID | Country | No. Case | No. Control |
|---|---|---|---|
| CRC1[4] | Austria | 46 | 63 |
| CRC2[5] | China | 73 | 54 |
| CRC3[6] | Germany | 60 | 60 |
| CRC4[7] | France | 53 | 61 |
| CRC5[8] | United States | 52 | 52 |

**Table S3**: Datasets of microbiome-metabolome association studies.

| Study ID | Correlated Sample Y/N | No. Subjects Control/Case | No. Samples Control/Case | Microbiome 16S/WGS | Metabolome Target/Untargeted |
|---|---|---|---|---|---|
| MTBL1[9] | N | 54/42 | 54/42 | WGS | Targeted |
| MTBL2[10] | N | 74*/220 | 74*/220 | WGS | Targeted |
| MTBL3[11] | N | 102/138 | 102/138 | 16S | Untargeted |
| MTBL4[12] | N | 56/164 | 56/164 | WGS | Untargeted |
| MTBL5[13] | Y | 24/51 | 139/305 | WGS | Targeted |
| MTBL6[14] | Y | 26/79 | 104/278 | WGS | Untargeted |
| MTBL7[15] | N | 67/220 | 67/220 | WGS | Untargeted |
| MTBL8[16] | Y | 83/NA | 164/NA | 16S | Untargeted |

*: the number is after removing control subjects shared with MTBL1.

**Tables S2 and S4 are too big and included in the Additional files 2 and 3.**
